# Single cell transcriptomics identifies adipose tissue CD271+ progenitors for enhanced angiogenesis in limb ischemia

**DOI:** 10.1101/2023.02.09.527726

**Authors:** Oto Inoue, Chiaki Goten, Daiki Hashimuko, Kosei Yamaguchi, Yusuke Takeda, Ayano Nomura, Hiroshi Ootsuji, Shinichiro Takashima, Kenji Iino, Hirofumi Takemura, Manasi Halurkar, Hee-Woong Lim, Vivian Hwa, Joan Sanchez-Gurmaches, Soichiro Usui, Masayuki Takamura

**Affiliations:** Department of Cardiovascular Medicine, Graduate School of Medical Sciences, Kanazawa University, Kanazawa, Japan; Department of Thoracic, Cardiovascular and General Surgery, Graduate School of Medical Sciences, Kanazawa University, Kanazawa, Japan; Division of Biomedical Informatics, Cincinnati Children’s Hospital Medical Center, Cincinnati, Ohio, USA; Department of Pediatrics, University of Cincinnati College of Medicine, Cincinnati, Ohio, USA; Division of Endocrinology, Cincinnati Children’s Hospital Medical Center, Cincinnati, Ohio, USA; Division of Developmental Biology, Cincinnati Children’s Hospital Medical Center, Cincinnati, Ohio, USA; Lead contact

**Author notes:** Correspondences (J.S.G.), (S.U.), (M.T.).

**Keywords:** single cell transcriptomics, therapeutic angiogenesis, cell therapy, CD271, adipose tissue, insulin resistance, peripheral artery disease, critical limb ischemia

## Abstract

Therapeutic angiogenesis using mesenchymal stem/stromal cell grafts have shown modest and controversial effects in preventing amputation for patients with critical limb ischemia. Through single-cell transcriptomic analysis of human tissues, we identified CD271^+^ progenitors specifically from subcutaneous adipose tissue (AT) as having the most prominent pro-angiogenic gene profile distinct from other stem cell populations. AT-CD271^+^ progenitors demonstrated robust *in vivo* angiogenic capacity, over conventional adipose stromal cell grafts, characterized by long-term engraftment, augmented tissue regeneration, and significant recovery of blood flow in a xenograft model of limb ischemia. Mechanistically, the angiogenic capacity of CD271^+^ progenitors is dependent on functional CD271 and mTOR signaling. Notably, the number and angiogenic capacity of CD271^+^ progenitors was strikingly reduced in insulin resistant donors. Our study highlights the identification of AT-CD271^+^ progenitors with *in vivo* superior efficacy for limb ischemia. Furthermore, we showcase comprehensive single-cell transcriptomics strategies for identification of suitable grafts for cell therapy.

**HIGHLIGHTS:**

- Adipose tissue stromal cells have a distinct angiogenic gene profile among human cell sources.
- CD271^+^ progenitors in adipose tissue have a prominent angiogenic gene profile.
- CD271^+^ progenitors show superior therapeutic capacities for limb ischemia.
- CD271^+^ progenitors are reduced and functionally impaired in insulin resistant donors.

**GRAPHICAL ABSTRACT:** 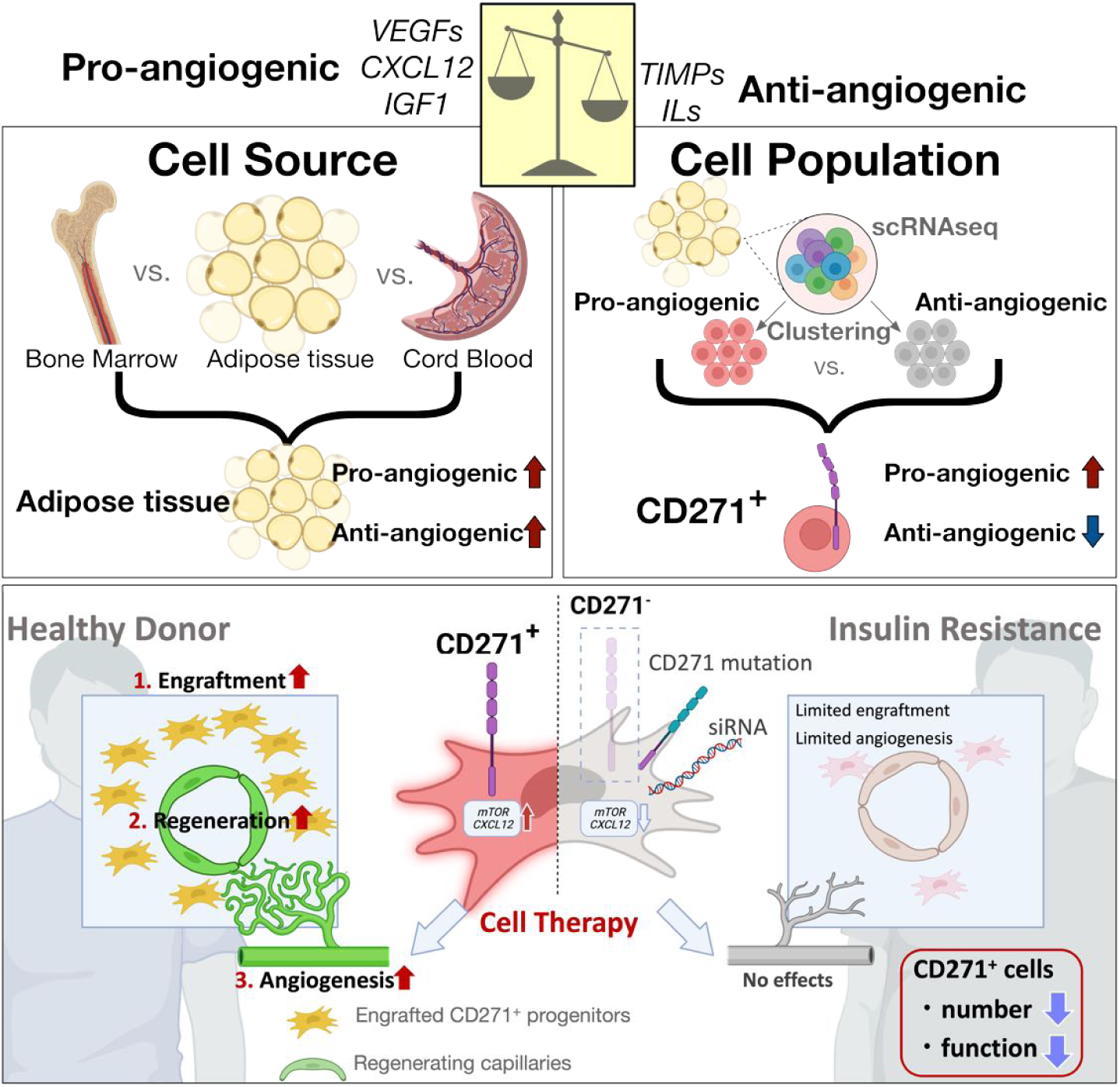

## INTRODUCTION

Critical limb ischemia (CLI), the end-stage of peripheral artery disease, is increasingly prevalent, in part due to the worldwide increase in obesity and diabetes.^1^ Prognosis for patients with CLI is poor, comparable to malignant diseases.^2^ One of the fatal features of CLI is atherosclerotic calcification, a condition difficult to treat even with currently available revascularization techniques such as catheter interventions and bypass surgery.^1^ Consequently, at least 1 in 3 CLI patients are not eligible for revascularization therapies.^3^ To combat this, the strategy of inducing neovascularization in ischemic tissues, known as therapeutic angiogenesis, has emerged as an attractive alternative therapy for CLI.^4^

Mesenchymal stem/stromal cells with regenerative capacities were discovered from multiple human organs including “harvestable” tissues such as bone marrow (BM), subcutaneous adipose tissue (AT), and umbilical cord blood (UCB).^5^ The main reported therapeutic mechanism associated with cell therapy-induced neovascularization is through secretion of growth factors (i.e., VEGF, HGF, SDF-1) and extracellular vesicles from injected cells. ^6, 7^ Preclinical studies of cell-based therapeutic angiogenesis using these cell sources showed favorable effects in animal models,^7–11^ but have translated poorly in clinical trials. ^4, 7^ For CLI, unsorted BM mononuclear cells are the most intensively evaluated in clinical trials. In a meta-analysis including 19 randomized controlled trials, cell therapy showed favorable effects on surrogate endpoints such as pain score, pain-free walking distance, and ankle brachial 65 index, but showed no significant benefits on primary endpoints including amputation-free rate and complete ulcer healing^12^. Prospective isolation of specific cell populations from BM mononuclear cells, based on CD133^+^ or CD34^+^ cells which were reported to be angiogenic preclinically,^10, 13^ also showed no evidence for lowering the risk of limb amputation in clinical trials.^14–16^ Thus, a suitable cell population(s) optimal for therapeutic angiogenesis to achieve limb salvation has yet to be identified.

Single-cell transcriptomic technologies has revolutionized the capacity to identify lineages in tissue homeostasis and markers of specific stem cell populations.^17–21^ However, application of single-cell transcriptomic analysis in cell therapy has been limited to only chimera antigen receptor T (CAR-T) cell therapy for lymphoma, where important cell populations for treatment response were assessed by single-cell RNA-sequencing (scRNA-seq). ^22, 23^

In this study, we took advantages of available scRNA-seq datasets generated from different human tissues to screen for cell surface markers that allow the segregation of a highly angiogenic subpopulation of cells. We identified that the CD271^+^ cells specifically from AT have a distinct angiogenic gene profile with promising *in vivo* angiogenic capacity. Mechanistically, CD271/mTOR axis showed an important role for *in vivo* angiogenic capacity. Furthermore, we extend our findings to evaluate the feasibility and revascularization efficacy of CD271^+^ cells from donors with cardiovascular disease (CVD). We obtained a sufficient number of CD271+ progenitors from metabolically healthy 83 CVD donors, and discovered superior therapeutic potentials over conventional cell grafts for patients suffering from CLI.

## RESULTS

### Adipose Tissue Lineage-negative Cells Show a Distinct Angiogenic Gene Profile among Human Stem Cell Sources

There is a strong need to identify the most suitable cell source for therapeutic angiogenesis. To search for subpopulations with pro-angiogenic capacity, we explored single cell gene expression profiles from readily available human stem cell sources. We analyzed available scRNA-seq datasets from bone marrow (BM),^24^ umbilical cord-blood (UCB),^25^ and stromal vascular fraction (SVF) of subcutaneous adipose tissue (AT)^26^ from healthy donors (***n*** = 2/tissue, **Figure 1A** and **Table S1**). We performed dimension reduction with t-distributed stochastic neighbor embedding (tSNE) and cell clustering of a total of 47,035 cells (**Figure 1A-B** **and Figure S1**). To annotate clusters, marker genes for each cluster were collated manually based on reported markers in the Human Cell Atlas and the Human Protein Atlas (proteinatlas.org, **Figure S1**). ^19, 27, 28^ We removed mature immune and endothelial cells, i.e. Lineage-positive (Lin^+^) cells,^27^ from further analysis to enrich stem/progenitor cell populations. Since classic lineage markers of immune and endothelial cells differ between tissues, and CD31^+^ cells are reported to be angiogenic only in BM^11^, we used a well-established but distinct set of lineage-markers in each of these tissues^29–31^ (listed in **Figure 1A**). Overall, we processed 18,053 Lin^-^ cells (BM; 9,289, UCB; 1,932, and AT; 6,832 cells, **Figure 1B**).

**Figure 1:**
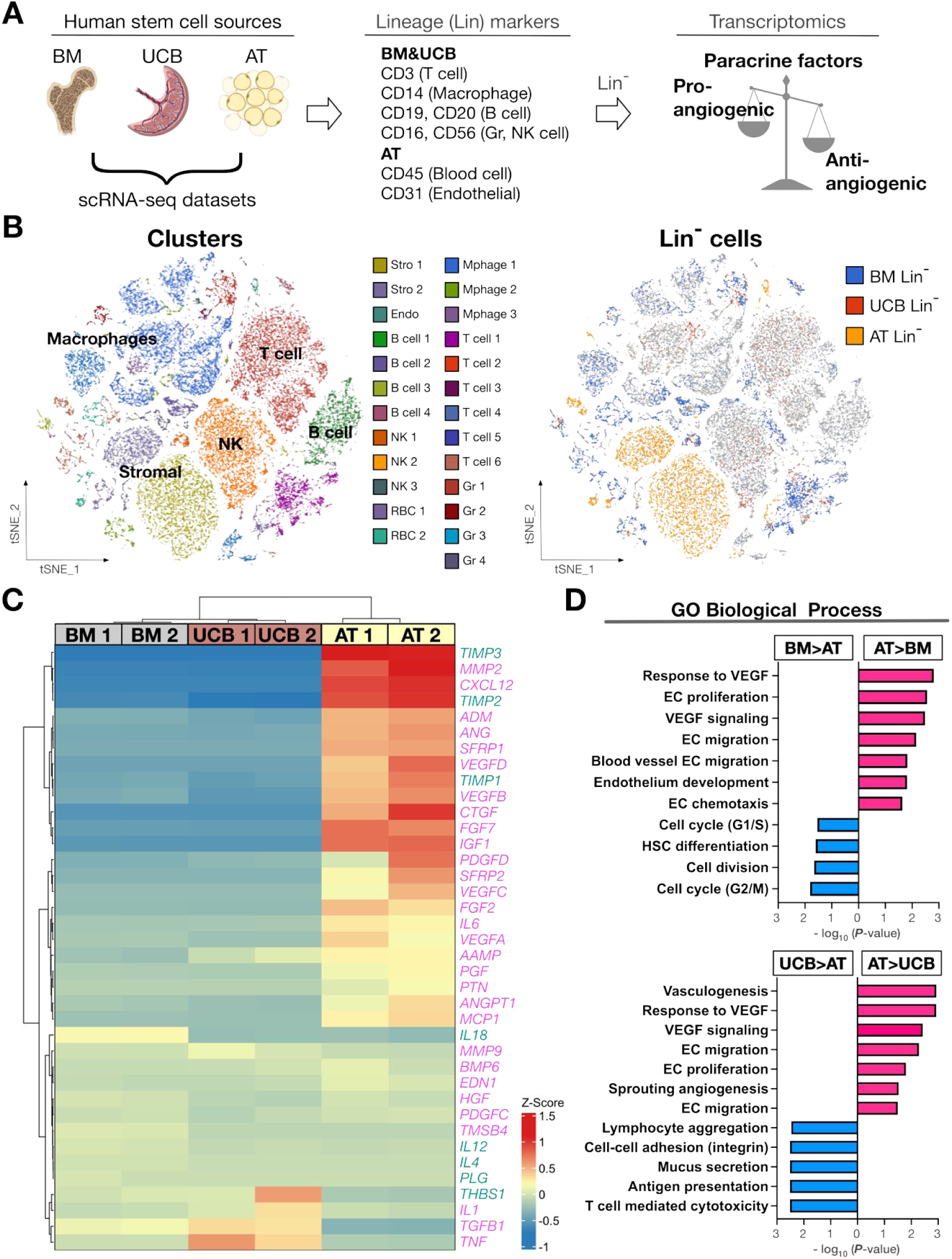
Adipose tissue shows an angiogenic gene profile in single-cell transcriptomics among human stem cell sources. (A) Workflow of scRNA-seq analysis to characterize pro/anti-angiogenic gene profiles of human cell sources. (B) tSNE map and clustering of 47,035 cells derived from BM, UCB, and AT (left). Distribution of BM, UCB, and AT-Lin^-^ cells (right). (C) Heatmap with hierarchical clustering of gene expressions of 38 pro-/anti-angiogenic growth factors in Lin^-^ cells from human BM, UCB, and AT. (D) Pathway enrichment analysis between indicated cell sources. The GO terms associated with angiogenesis (red) or overrepresented on BM/UCB (blue) are displayed. See also **Figure S1**.

The main mechanism of cell-based therapeutic angiogenesis has been reported to be paracrine activities of soluble growth factors released by injected cells ^6, 32^. Thus, to further characterize angiogenic gene profiles of stem cell sources, we used a gene set of reported pro/anti-angiogenic factors released from tissue stem cells^6^. We updated the list with an additional gene set of 38 well-known secreted pro-/anti-angiogenic factors ^33–35^ (**Table S2**). Hierarchical clustering of BM, UCB, and AT Lin^-^ cells based on expression of pro-/anti-angiogenic factor genes clearly isolates AT cells as having a distinctive gene expression profile. This was characterized by prominent expression of pro-angiogenic factors including *MMP2*, *CXCL12*, *VEGFs* and *IGF1* (**Figure 1C**). Consistently, gene enrichment analysis resulted in a number of pro-angiogenic pathways which were significantly enriched in AT compared to BM or UCB (**Figure 1D**). On the other hand, we observed that anti-angiogenic factors such as *TIMP2* and *TIMP3* were also upregulated in AT (**Figure 1C**). Therefore, we hypothesized that AT would be an attractive cell source if pro-angiogenic cells could be segregated from TIMP2/3 expressing anti-angiogenic cells.

### Adipose Tissue scRNA-seq Analysis Filtered 30 Surface Markers of Angiogenic Cells

To delineate angiogenic populations within the AT, we analyzed Lin^-^CD34^+^ progenitors on available scRNA-seq datasets from six non-diabetic human subcutaneous AT SVF (**Figure 2A**). Louvain clustering divided AT Lin^-^CD34^+^ progenitor cells^8, 36^ into 3 progenitor groups (**Figure 2B** **and Figure S2A**). To understand the nature of each of these 121 clusters, we analyzed the top 10 marker genes of each Progenitor cluster (**Figure 2C**). Progenitor 1 cluster was characterized by higher expression of adipocyte lineage markers (*APOE,* and *CEBPD*), while Progenitor 2 by mesenchymal/fibrotic markers (*FBN1*, *MFAP5*, and *FN1*). Progenitor 3 highly expressed some adipocyte lineage markers common to Progenitor 1, including *APOD*, but was characterized by higher expression of non-adipocyte markers *MYOC* and *IGFBP3*. Among these 3 clusters, pathway enrichment analysis found that Progenitors 1 & 3 were dominated by genes involved in sprouting angiogenesis and lipogenesis (**Figure 2D**). Based on this analysis, Progenitors 1 & 3 were defined as pro-angiogenic clusters while Progenitor 2 was defined as an anti-angiogenic cluster. Using the same gene set as for cell source screening (**Figure 1C** and **Table S1**), we compared pro/anti-angiogenic factor genes between the pro- and anti-angiogenic clusters. The expression level of pro/anti-angiogenic factors clearly separates pro- and anti-angiogenic clusters on our reanalyzed scRNA-seq datasets from six non-diabetic human subcutaneous AT SVF (**Figure 2A, S2B**). Among the 38 genes in the gene set, 22 genes were differentially expressed between pro/anti-angiogenic clusters. As expected, the pro-angiogenic cluster showed higher expression of 12 pro-angiogenic factors, while all 5 anti-angiogenic factor genes were highly expressed in the anti-angiogenic cluster (**Figure S2B**).

**Figure 2:**
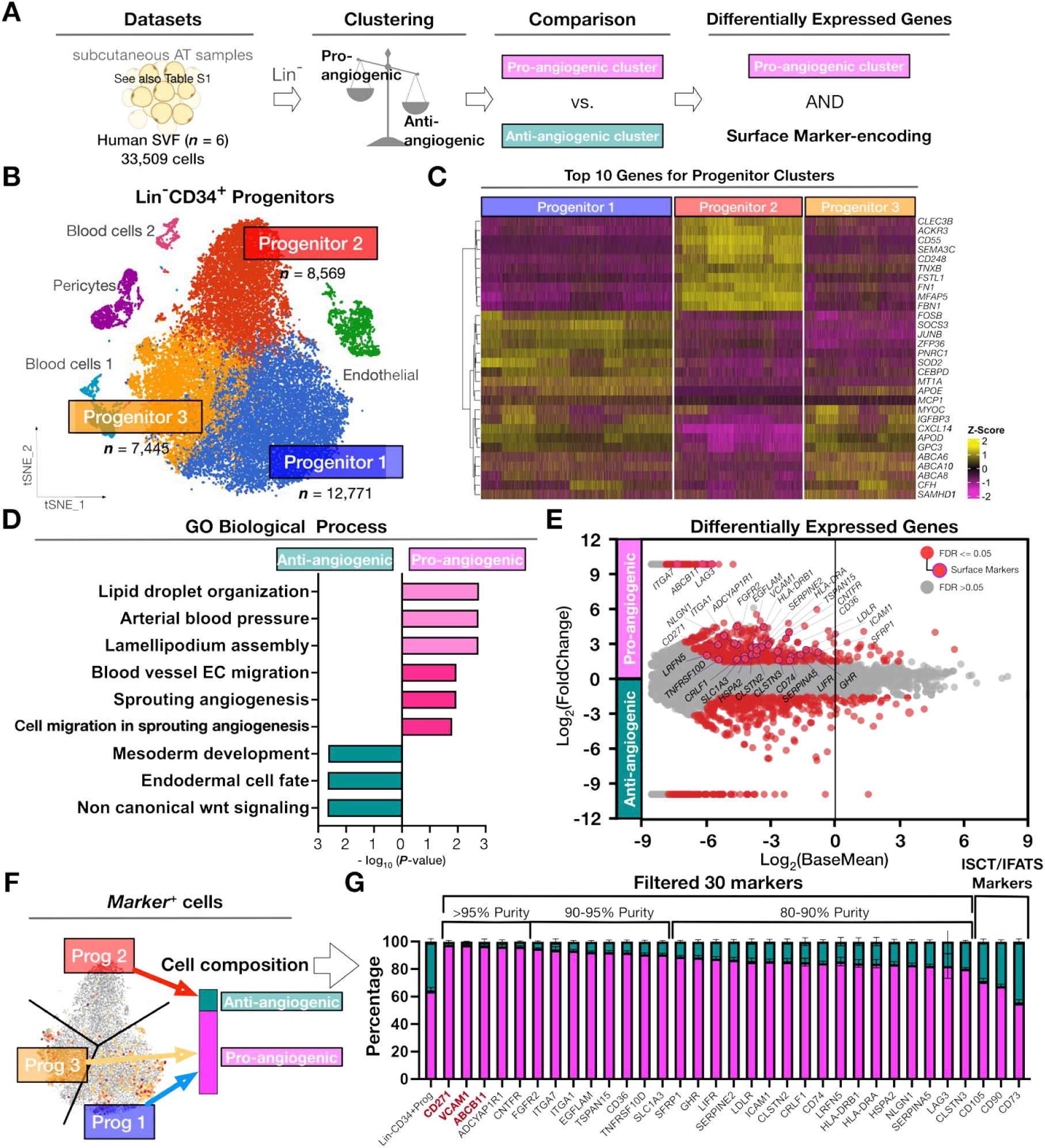
Adipose tissue single-cell transcriptomics filters 30 surface markers of angiogenic cluster cells. (A) Diagram illustrating filtering strategy of surface marker genes specific for pro-angiogenic cluster in human SVF. (B) tSNE map and clustering of 33,509 SVF cells obtained from 6 human subcutaneous AT. (C) Single-cell heatmap of top 10 marker gens among 3 progenitor clusters (***n*** = 6). (C) Pathway enrichment analysis between pro-/anti-angiogenic clusters. The top 3 up/downregulated and angiogenesis-related GO terms are displayed (***n*** = 6). (D) Volcano plot of 2,621 differentially expressed genes between pro/anti-angiogenic clusters. Red dots show differentially expressed genes with |log2(FoldChange)| > 1.5. Violet circles highlight genes encoding surface proteins (***n*** = 6). (E) Diagram illustrating the ranking analysis of 30 surface marker genes based on the purity of pro-angiogenic cluster. (F) Distribution of the pro/anti-angiogenic clusters within the surface marker expressing cell populations (***n*** = 6). Data are presented as the mean ± SEM. See also **Figure S2**.

To obtain intact cell populations for cell therapy purposes, purification procedures utilizing surface markers are necessary. Thus, we next focused on identifying 139 the surface markers specifically expressed in the pro-angiogenic cluster that would be suitable for downstream flow cytometric isolation. Based on 2,621 differentially expressed genes between pro-/anti-angiogenic clusters from scRNA-seq analysis identified when grouping data from the six samples analyzed (**Figure 2E**, **Figure S2C and Table S3**), a total of 30 surface protein-encoding genes (**Table S4**) were identified. We concluded that these 30 surface proteins may potentially lead to the specific isolation of pro-angiogenic cells.

Next, we determined which marker from the filtered 30 surface protein-encoding genes would be the most suitable for cell graft preparation for therapeutic angiogenesis. To achieve this, we evaluated the specificity of the 30 individual surface proteins enriched in the pro-angiogenic cluster by analyzing the number of cells that express each protein in each cluster using the data from the same six non-diabetic human subcutaneous AT SVF (**Figure 2A, 2F**).

From this analysis, 5 surface markers were identified for isolating pro-angiogenic cells with over 95% purity (**Figure 2G**). Among them, CD271 (Low-affinitive nerve growth factor receptor, LNGFR) emerged as the top-ranked surface marker, closely followed by VCAM1 (vascular cell adhesion molecule 1) and ABCB11 (ATP binding cassette subfamily B member 11). In particular, from the cells detected to express CD271, 98% were on the pro-angiogenic cluster (**Figure 2G**). This is appreciably higher compared to Lin^-^CD34^+^ progenitors (Prog 1,2, and 3) and to other previously known AT progenitor markers37 such a 157 s CD105, CD90, and CD73 (**Figure 2G**). Furthermore, out of the cells labelled by CD271 expression less than 3% were on the anti-angiogenic cluster, which was the lowest contamination rate amongst all studied markers (**Figure 2G**). Thus, we identified surface markers that may efficiently select for pro-angiogenic cells.

### Adipose Tissue CD271^+^ Cells are Highly Angiogenic

Next, we further analyzed the top three surface markers identified in our analysis aiming to find which cell population showed the highest angiogenic potential. We noted that CD271^+^, VCAM1^+^, and ABCB11^+^ cell subpopulations are distinct, with very few cells carrying all three markers, or even 2 of the 3 markers (**Figure 3A**). In particular, over 80% of CD271^+^ cells were VCAM^-^/ABCB11^-^. Therefore, we performed pathway enrichment analysis for subpopulations carrying each single marker, i.e. comparing between CD271^+^VCAM^-^ABCB11^-^, CD271^-^ VCAM^+^ABCB11^-^, and CD271^-^VCAM^-^ABCB11^+^. Strikingly, CD271^+^ cells showed pro-angiogenic pathway enrichment over VCAM1^+^ or ABCB11^+^ cell populations (**Figure 3B**). Direct comparison between CD271^+^ and CD271^-^ populations showed that CD271^+^ cells have higher expression of important pro-angiogenic factors (e.g., *MCP1*, *CXCL12*, *CTGF*, and VEGFA **Figure 3C**). Additionally, anti-angiogenic factors *TIMP2* and *TIMP3* were downregulated in CD271^+^ cells compared to CD271^-^ cells (**Figure 3C**).

**Figure 3:**
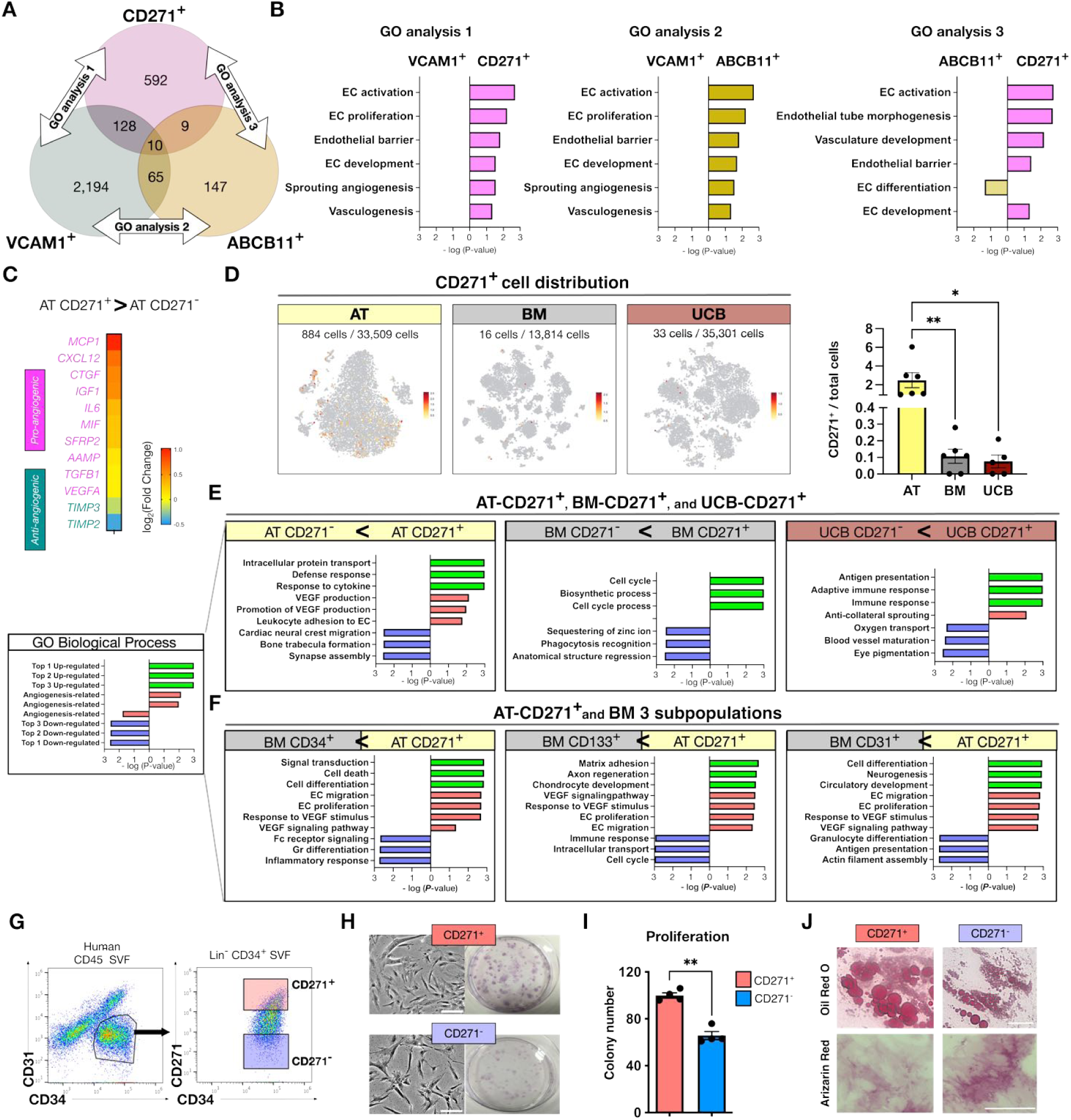
Adipose tissue CD271^+^ cell population is highly angiogenic. (A) Venn diagram illustrating the overlap between CD271^+^, VCAM1^+^, and ABCB11^+^ cells in Lin^-^CD34^+^ progenitor cells in AT scRNAseq analysis (***n*** = 6). (B) Pathway enrichment analysis between CD271^+^, VCAM1^+^, and ABCB11^+^ cell populations. (C) Heatmap of representative differentially expressed genes of pro-/anti-angiogenic factors between CD271^+^ and CD271^-^ cells. The color shows the value of log2 Fold Change. (***n*** = 6). (D) The distribution of CD271^+^ cells in tSNE map and quantification of the frequency of CD271^+^ cells in AT, BM, and UCB (***n*** = 2/organ). The color of dot plots represents the expression level of CD271. (E) Pathway enrichment analysis between CD271^+^ and CD271^-^ cells in AT, BM, and UCB. The top 3 up/downregulated and angiogenesis-associated GO terms are displayed (***n*** = 2/organ). (F) Pathway enrichment analysis between AT-CD271^+^ cells and three BM subpopulations. The top 3 up/downregulated and angiogenesis-associated GO terms are displayed (***n*** = 2/organ). (G) Representative flow cytometric plots from human subcutaneous AT SVF. (H) Representative images of cultured ADSCs and generated colonies (Giemsa staining) in CFU assay. Bars = 50 μm. (I) Colony number in CFU assay of CD271^+^ and CD271^-^ ADSCs (***n*** = 4/group). (J) Representative images of differentiation assays of CD271^+^ and CD271^-^ cells. Bars = 100 μm. Data are presented as the mean ± SEM. **\**P*** < 0.05 and \*\****P*** < 0.01.

We next asked if it is possible to obtain sufficient number of CD271^+^ cells from AT for therapeutic angiogenesis. We found that the number of CD271 expressing c 175 ells were at least 10 times higher in AT than in BM or UCB (**Figure 3D**). To confirm these results, we performed flow analysis on SVF cells of human subcutaneous AT. Consistently, we detected CD271 protein in at least 5–10% of total SVF cells of human subcutaneous AT (**Figure 3G**); in contrast, it was previously reported that less than 0.1% of individual BM cells expressed CD271 protein by flow analysis.^38^ From these data, we concluded that AT is the most suitable tissue for harvesting sufficient number of CD271^+^ cells.

Although AT carried higher numbers of CD271^+^ cells than either BM or UCB, we assessed whether CD271^+^ cells are pro-angiogenic, independent of tissue source. As expected, AT-CD271^+^ cells showed strong enrichment in proangiogenic pathways when compared to AT-CD271^-^ cells (**Figure 3E**). BM-CD271^+^ cells, in contrast, were characterized by pathways related to the cell cycle and proliferation (**Figure 3E**), consistent with a previous report that BM-CD271^+^ cells possess clonogenic and multi-lineage differentiation capacities.^38^ On the other hand, UCB-CD271^+^ cells showed upregulation of immune system-related pathways when compared to UBC-CD271^-^ (**Figure 3E**). These transcriptomic analyses indicated that although CD271^+^ cells are available from different tissues, AT-CD271^+^ cells exhibit a distinct, pro-angiogenic, gene profile.

It is of note that several subpopulations of BM, including CD34^+^ ^14^, CD133^+^ ^16^, and CD31+ 11 populations, have been reported to be angiogenic and also 193 have been tested in clinical trials. Thus, we compared transcriptomic profile of AT-CD271^+^ cells to these three BM subpopulations (**Figure 3F**). We found that AT-CD271^+^ cells consistently showed enrichment of pathways related to cell differentiation while BM subpopulations were characterized by involvement with immune systems. More importantly, AT-CD271^+^ cells demonstrated enrichment of pro-angiogenic pathways (e.g. activating endothelial proliferation and VEGF signaling) when compared to each BM subpopulation. Altogether, these transcriptomic data indicate that AT-CD271^+^ cells has higher angiogenic potential than candidates already in clinical trials.

### AT-CD271^+^ progenitors are culture-expandable with classical mesenchymal stem cell capacity

It is known that culture-expandable mesenchymal stem cell-like cells can be generated from unsorted AT-progenitors, and are commonly referred as adipose-derived stem cells (ADSCs) ^39^. To better understand human AT-CD271^+^ cell biology, we sorted and established *in vitro* cultures of CD271^+^ and CD271^-^ cells from subcutaneous AT of human healthy subjects (**Figure 3G**). Both types of cells were adherent and proliferative with similar spindle-shaped morphology (**Figure 3H**). CD271^+^ cells, however, showed enhanced proliferative capacity compared to CD271^-^ cells as determined by colony formation assay where CD271^+^ cells generated larger and more dense colonies (**Figure 3H**) in significantly higher number (**Figure 3I**). Both cell types readily differentiated into adipocyte (Oil Red O 211) and osteoblast (Alizarin Red) lineages (**Figure 3J**). Collectively, these *in vitro* assays characterized CD271^+^ progenitors as culture-expandable with classical ADSC morphology and capacities.^39^

### CD271^+^Progenitors Demonstrate Enhanced In Vivo Angiogenic Capacity with Long-Term Engraftment

We next examined the *in vivo* angiogenic capacity of CD271^+^ progenitors in a xenograft model of limb ischemia (**Figure 4A**). For this, we injected PKH-labeled human CD271^+^ or CD271^-^ progenitors (0.5 million cells / mice) directly into limb muscles of immunocompromised mice that are subjected to left femoral artery ligation. Cell-treated limbs were harvested at day 7, 14, and 35 after fluorescence-conjugated isolectin perfusion to detect capillaries with blood flow^9,30,40^.

**Figure 4:**
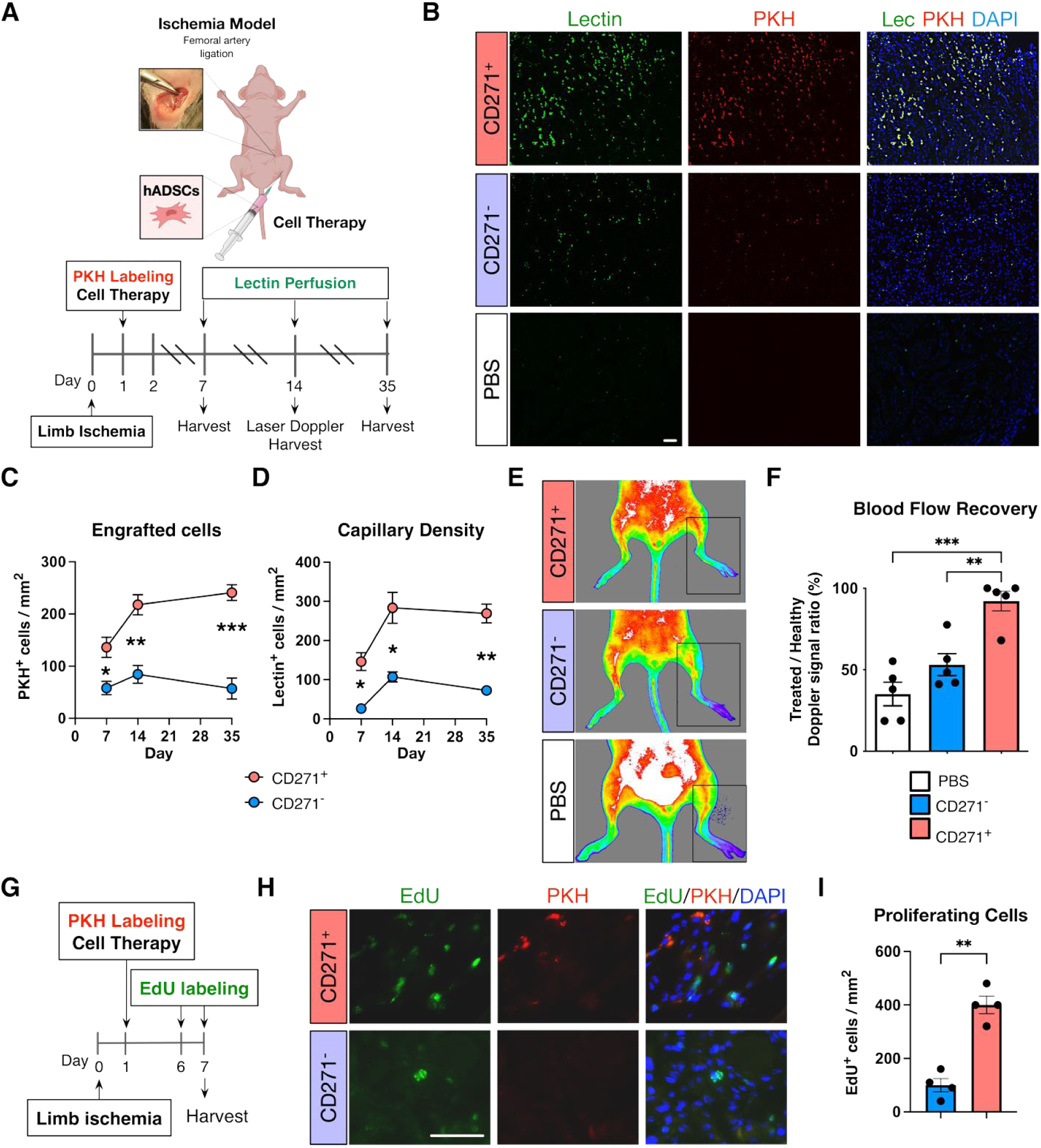
AT-CD271^+^ progenitors have high *in vivo* angiogenic capacity with long-term engraftment. (A) Diagram of our cell therapy protocol using PKH-labeled human ADSCs and fluorescence-conjugated lectin perfusion. (B) Representative IHC images for lectin^+^ capillaries and PKH^+^ engrafted human ADSCs at day 14 after cell transplantation with either CD271^+^ or CD271^-^ progenitors. Bar = 100 μm. (C) Quantitative analysis of engrafted ADSCs in ischemic limbs on day 7, 14, and 35 with either CD271^+^ or CD271^-^ progenitors (***n*** = 5). (D) Quantitative analysis of capillary density in ischemic limbs on day 7, 14, and 35 (***n*** = 5). (E) Representative images of laser Doppler analysis at day 14 after cell therapy. (F) Quantitative analysis of blood flow using laser Doppler analysis at day 14 after cell therapy with either CD271^+^ or CD271^-^ progenitors. Signal of treated limb is normalized to signal from healthy limb (set as 100%) (***n*** = 5). (G) Diagram of cell therapy experiments to evaluate in vivo proliferating cells in ischemic limb after cell therapy. (H) Representative IHC images for EdU^+^ cell nuclei and PKH^+^ engrafted human CD271^+^ or CD271^-^ progenitors at day 7 after cell therapy. Bar = 50 μm. (I) Quantitative analysis of EdU^+^ proliferating cells after CD271^+^ or CD271^-^ progenitor treatment (***n*** = 4). Data are presented as the mean ± SEM. \****P*** < 0.05, \*\****P*** < 0.01 and \*\*\****P*** < 0.005. See also **Figure S3**.

In this xenograft model, we observed that CD271^+^ progenitors clearly engrafted better than CD271^-^ progenitors in ischemic limb, with 2-3 times higher PKH^+^ cell numbers (**Figure 4B-C**). The enhanced differences in engraftment capacity of AT-CD271^+^ progenitors persisted for up to 5 weeks. Further, immunofluorescent analysis of tissue sections shows that AT-CD271^+^ progenitors localize around lectin^+^ capillaries (**Figure S3A**). To clarify whether injected CD271^+^ progenitors were potentially trans-differentiating to endothelial cells, we performed flow cytometric analysis of PKH^+^ cells from digested limb muscles 14 days after cell engraftment (**Figure S3B**). We confirmed that only 6% of engrafted 229 CD271+ progenitors expressed the endothelial marker CD31, which suggested the majority of the injected cells did not undergo differentiation into mature endothelial cells, at least not within the time frame of the protocol (**Figure S3C**). On the other hand, isolated PKH^+^ cells maintained expression of human CD271 and typical markers of cultured ADSCs^37^ (human CD90 and CD105) (**Figure S3C**). These findings confirmed that engrafted CD271^+^ progenitors preserve their stem cell marker expressions and likely acted as stem/progenitor cells to support intrinsic regeneration.

Correlating with higher survival of engraftment, CD271^+^ progenitor therapy induced enhanced revascularization after limb ischemia. This is evidenced by 3-fold higher capillary density (Lectin^+^ and mouse CD31^+^, **Figure S3D**) than that induced by CD271^-^ progenitors (**Figure 4D**). Strikingly, CD271^+^ progenitors successfully led to recovery of blood flow in treated limb, to a level comparable to healthy limbs, as determined by both imaging and doppler blood flow analysis (**Figure 4E-F**).

To confirm tissue reparative capacity of CD271^+^ progenitors, we incorporated *in vivo* EdU labeling into our *in vivo* protocol (**Figure 4G**). Since neovascularization and blood flow recovery were favorably accomplished by day 14 in our model, we evaluated EdU incorporation on day 7 when tissue neovascularization was in progress. *In vivo* EdU labeling demonstrated that treatment with CD271^+^ progenitors led to approximately four times higher proliferation of host cells (largely composed of skeletal muscle cells and endothelial 247 cells and identified as PKH-) than treatment with CD271^-^ progenitors (**Figure 4H-I**). Collectively, the data from our xenograft model provide promising translational *in vivo* angiogenic capacity of CD271^+^ progenitors with long-term engraftment and enhanced host cell proliferation.

### Mutant CD271 Impairs the Angiogenic Gene Profile and Therapeutic Capacity of mouse ADSCs

We next explored potential mechanisms that drove angiogenic capacity of CD271^+^ progenitors. Reanalysis of scRNA-seq datasets of SVF cells from neonatal mice inguinal fat pads^41^ subclustered cells into 10 clusters (**Figure 5A** **and Figure S4A-B**). CD271^+^ progenitors were distributed predominantly with Schwann cell, pericytes, and Lin^-^ Progenitor clusters (**Figure S4B**). Consistent with human CD271^+^ progenitors gene expression profiles (**Figure 3C**), mouse CD271^+^ showed upregulation of pro-angiogenic factors including *Igf1* and *Cxcl12*, and downregulation of anti-angiogenic factors *Timp2* and *Timp3*. (**Figure 5B**). Furthermore, mouse AT-CD271^+^ progenitors showed enrichment of angiogenesis-related GO terms (endothelial cell proliferation and VEGF signaling pathway, **Figure 5C**). We asked whether a functional CD271 protein is necessary for the angiogenic capacity of CD271^+^ progenitors in mice. We used mice lacking exon 3 of the CD271 gene which encodes three of the four necessary cysteine-rich repeats in the extracellular domain. The loss of exon 3 generated an in-frame mutant CD271 protein with a 2-to 3-fold reduction in ligand binding42 (**Figure 265 5D**). We established cultured mouse ADSCs from wild-type littermates (Control) and CD271 mutant mice (CD271^Mut^). Compared to Control-ADSCs, CD271^Mut^ ADSC transplants demonstrated impaired cell engraftment (**Figure 5E-F**), impaired *in vivo* angiogenesis (**Figure 5G**), and reduced blood flow recovery (**Figure 5H-I**). This data suggests that normal CD271 activity is required for CD271^+^ cells to have full angiogenic capacity. This highlights an important new biological function of CD271 and proposes the concept that CD271 is not only a surface marker for cell selection.

**Figure 5:**
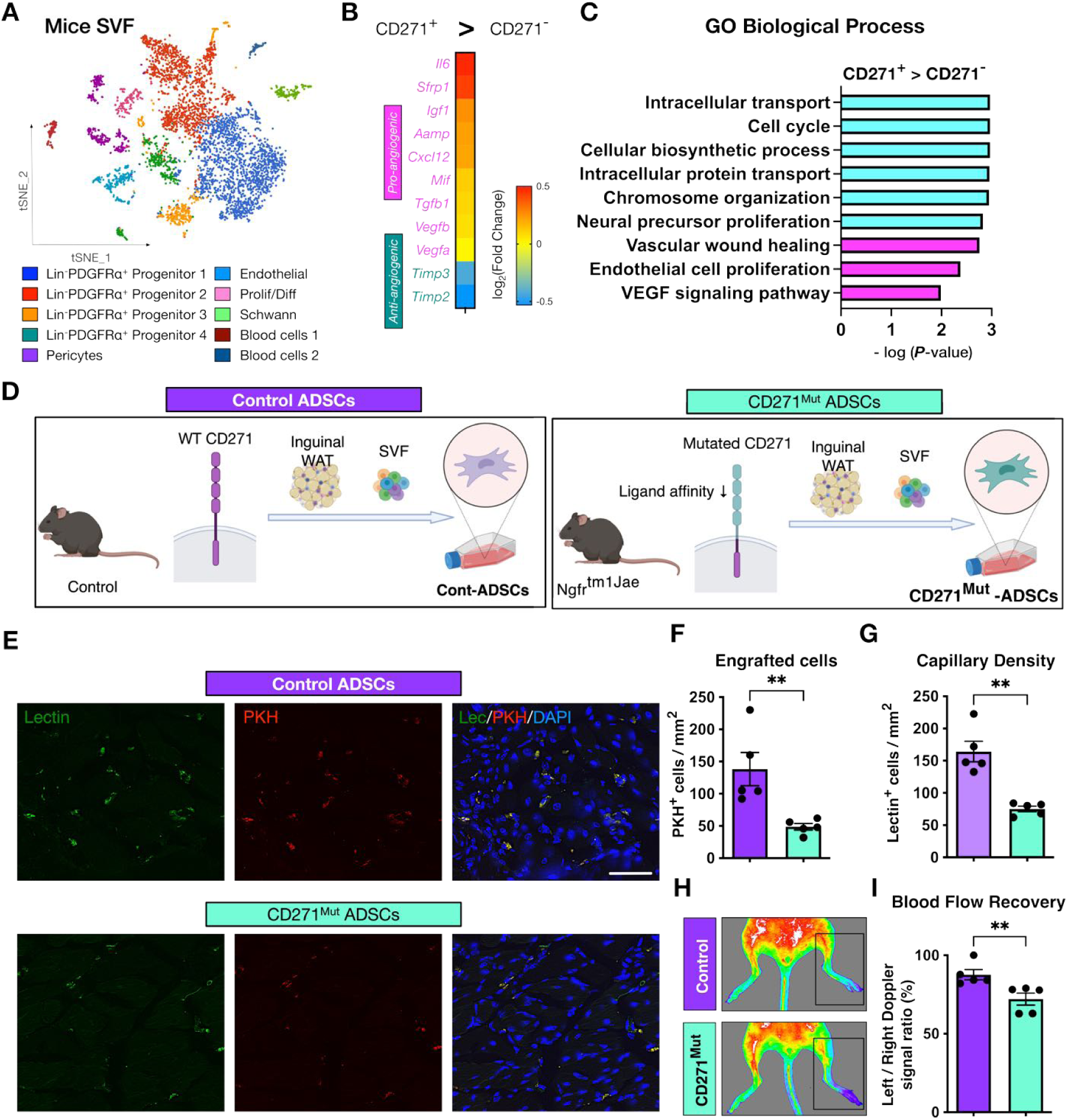
Normal CD271 activity is required for proangiogenic gene profile and *in vivo* angiogenic capacity of mouse ADSCs. (A) tSNE map and clustering of 4,912 neonatal mouse inguinal fat pad SVF cells. (B) Heatmap presentation of representative differentially expressed genes of pro/anti-angiogenic factors between CD271^+^ and CD271^-^ mouse cells. (C) Pathway enrichment analysis between mouse CD271^+^ and CD271^-^ mouse cells in Lin^-^ PDGFRα^+^ progenitors. The top 6 upregulated and angiogenesis-associated GO terms are displayed. (D) Diagram illustrating preparation of CD271-mutated mouse-derived (CD271^Mut^) and littermate wild-type mouse-derived (Control) ADSCs. (E) Representative IHC images of lectin^+^ capillaries and PKH^+^ engrafted mouse ADSCs on day 14 after cell transplantation with either CD271^Mut^ or control ADSCs. Bar = 50 μm. (F) Quantitative analysis of engrafted ADSCs in ischemic limbs on day 14 after treatment with CD271^Mut^ or control ADSCs. (***n***=5) (G) Quantitative analysis of capillary density in ischemic limbs on day 14 after treatment with CD271^Mut^ or control ADSCs. (***n***=5) (H) Representative images of laser Doppler analysis on day 14 after transplantation with CD271^Mut^ or control ADSCs following limb ischemia induction (***n*** = 5). (I) Quantitative analysis of blood flow using laser Doppler analysis at day 14 after cell therapy with either CD271^Mut^ or control ADSCs. Signal of treated limb is normalized to signal from healthy limb (set as 100%). Data are presented as the mean ± SEM. \****P*** < 0.05 and ***\*\*P*** < 0.01. See also **Figure S4**.

We asked what downstream signaling pathways of CD271 may participate in the angiogenic functions of CD271^+^ progenitors. CD271 can induce cell death or cell survival, depending on interacting coreceptors.^43^ Among the three known coreceptors of CD271, tropomyosin receptor kinase B (TRKB), which coordinates with the mTOR pathway for cell survival^44^ (**Figure S5A**), was predominantly expressed in human AT-CD271^+^ progenitors while cell expressing the other coreceptors were rarely found (**Figure S5B**). We confirmed that over 95% of CD271^+^ progenitors co-expressed TRKB protein even under cultured condition (**Figure S5C**). Cultured CD271^+^ progenitors also showed higher expression of *MTOR* than CD271^-^ progenitors (**Figure S5D**). Furthermore, the known downstream genes regulated by mTOR signaling, i.e., *HIF1A* and *VEGFA*, were significantly upregulated in CD271^+^ progenitors compared to CD271^-^ (**Figure S5D**). Consistent with previous reports of positive feedback between mTOR and SDF-1 (CXCL12)45, *CXCL12* 283 was also highly expressed in CD271+ progenitors (**Figure S5D**). Interestingly, we noted that in mouse CD271^Mut^ ADSCs, *Mtor* expression was reduced by ∼50% (**Figure S5E**), which resulted in concomitant inhibition of downstream factors *Hif1a*, *Vegfa*, and *Cxcl12* (**Figure S5E**). From these observations, we hypothesized that the downstream mTOR pathway activated by CD271 may be required for the angiogenic capacity of CD271^+^ progenitors. To test this hypothesis, we knockdown mTOR in human ADSCs (cultured unsorted AT cells) using short interfering RNAs (si-mTOR ADSCs). Gene expression analysis showed efficient knockdown of mTOR (**Figure S5F**). Additionally, si-mTOR ADSCs shown downregulated expression of *VEGFA* and *CXCL12* (**Figure S5F**). Next, we functionally tested the effects of decreased mTOR levels in cell therapy experiments using our ischemic limb mouse model. si-mTOR treatment resulted in loss of *in vivo* angiogenic capacity of ADSCs as evaluated by capillary density (**Figure S5G-H**), engraftment (**Figure S5I**) and tissue reparative capacity evaluated by the number of proliferating cells (**Figure S5J-K**). Together, our findings strongly suggest that a functional CD271/mTOR axis is necessary for full ADSC angiogenic capacities.

### CD271^+^ Progenitors from Insulin Resistant Patients are Decreased and Have Poor Therapeutic Potential

Next, we asked if AT-CD271^+^ progenitors are available in sufficient number to be used in clinical settings. For this, we analyzed patient-derived AT samples. F 301 irst, we evaluated the number of CD271^+^ progenitors in subcutaneous AT from 38 CVD patients with various pathological conditions and diseases (**Table S4**). The frequency of CD271^+^ progenitors in SVF cells (CD271^+^/SVF) ranged from 0.2 to 40% and was not significantly affected by donor sex, BMI, or age (**Figure 6A**). This is clearly favorable for cell preparation from a variety of donors. Intriguingly, the number of CD271^+^ progenitors was negatively correlated with the insulin resistance score (HOMA-IR) (**Figure 6A**). Consistently, CD271^+^ progenitor number was reduced by approximately 75% in patients with insulin resistance (HOMA-IR > 2.9)^46^ as determined by flow cytometry (**Figure 6B**). Because immune cells accumulate in insulin-resistant AT, we additionally calculated the percentage of CD271^+^ progenitors in Lin^-^CD34^+^ cells. Consistently, CD271^+^/Lin^-^CD34^+^ SVF was also significantly reduced in patients with insulin resistance (**Figure 6B**). To validate the finding of our 38 CVD patient cohort with an orthogonal method, we analyzed available subcutaneous AT scRNA-seq datasets from lean healthy (insulin sensitive) subjects versus obese type 2 diabetic (insulin resistant) subjects (**Figure 6C** **and Figure S6A**). We found a significant reduction in the number of CD271^-^ expressing cells in AT from insulin resistant subjects (**Figure 6D-E**). Furthermore, GO enrichment analysis shows decreased angiogenesis-related terms in CD271^+^ progenitors from insulin resistant donors when compared to CD271^+^ progenitors from insulin sensitive donors (**Figure 6F** **and Table S5**).

**Figure 6:**
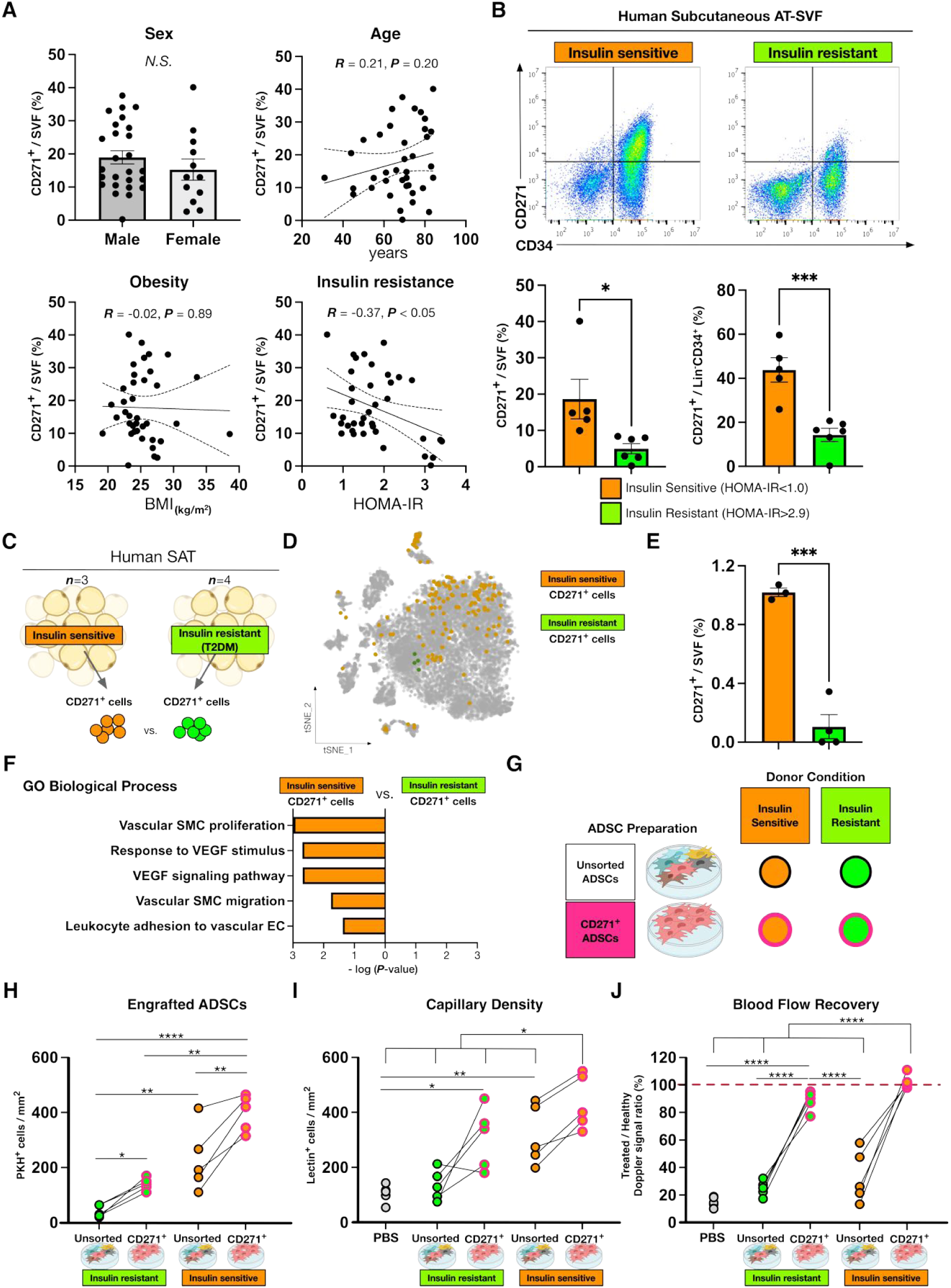
AT-CD271^+^ progenitors are decreased and impaired in insulin resistant donors. (A) Plots comparing CD271^+^ cell frequency in SVF (CD271^+^/SVF) and clinical parameters of CVD patients with or without insulin resistance (***n*** = 38). (B) Representative flow cytometric plots and comparative analysis of CD271^+^ cell abundance in total SVF or Lin^-^CD34^+^ SVF of CVD patients with or without insulin resistance (***n*** = 5-6). (C) Diagram illustrating human AT samples used for scRNA-seq analysis (D) Distribution of CD271^+^ cells in tSNE map derived from lean healthy (***n*** = 3, insulin sensitive, **blue dots**) and obese-diabetic patients (***n*** = 4, insulin resistant, **red dots**). (E) Comparative analysis of the percentage of CD271^+^ cells between patients with and without insulin resistance in the scRNA-seq analysis (***n*** = 3-4). (F) Pathway enrichment analysis between CD271^+^ cells from patients with and without insulin resistance. GO terms associated with angiogenesis are displayed. (G) Diagram illustrating ADSC groups for cell therapy experiments stratified by donor types and ADSC preparation methods (unsorted and CD271^+^ ADSCs). (H) Quantitative analysis of engrafted ADSCs and capillary density in ischemic limbs at day 14 after cell therapy (***n*** = 5). The red-colored dashed line indicates complete recovery of blood flow (100%). \****P*** < 0.05, \*\****P*** < 0.01, \*\*\****P*** < 0.005, and \*\*\*\****P*** < 0.001. See also **Figure S6**.

Since our results show that CD271^+^ progenitors have a positive therapeutic effect in our mouse ischemic limb model, we next hypothesized that CD271^+^ progenitor therapy is more beneficial over conventional unsorted ADSCs in our model. Hence, we prepared paired unsorted ADSCs and sorted CD271^+^ progenitors from 5 insulin resistant and 5 insulin sensitive patient subgroups from our 38-patient cohort (**Figure 6A-B**) to test their therapeutic effectiveness in our ischemic limb mice model (**Figure 6G**). As shown in **Figure 6H**, sorted human CD271^+^ progenitors showed markedly enhanced engraftment over conventional ADSCs from the same donors. Notably, cell therapy with CD271^+^ progenitors of metabolically healthy donors resulted in improved engraftment than CD271^+^ progenitors of insulin resistant donors (**Figure 6H**). Interestingly, although CD271^+^ progenitors from both insulin resistant and metabolically healthy donors increased capillary density compared to respective unsorted cells, CD271^+^ progenitors of metabolically healthy donors achieved striking higher capillary density than those from insulin resistant donors (**Figure 6I**). It is important to note that conventional unsorted ADSCs from insulin resistant donors showed capillary density equivalent to PBS injection suggesting very limited effect of this population (**Figure 6I**). Finally, CD271^+^ progenitors from insulin sensitive donors achieved full recovery of blood flow (**Figure 6J**) while CD271^+^ progenitors from insulin resistant donors were less effective but still achieved a remarkable > 85% recovery. These data further highlight the promising efficacy of CD271^+^ progenitor therapy for ischemic limb, even from insulin resistant donors, albeit therapy from insulin s 337 ensitive donors may lead to better long-term outcomes as engraftment is higher and improve the angiogenic capacity of the ischemic lesion.

### Preliminary Positive Impacts of Autologous CD271^+^ Progenitors in a Trial Outcome of SVF Therapy for CLI

To further investigate the therapeutic potential of CD271^+^ progenitors, we retrospectively evaluated data from a limited human cell therapy trial (multi-center clinical trial of SVF therapy) for no-option CLI patients^47^ (**Figure 7A** and **7B**). During the one-year follow-up period, patients received no interventional treatments (e.g. catheter intervention, surgery). Medications were already optimized prior to SVF therapy under consensus guidelines, and no further changes were made ^48^. Of the 4 patients within our trial who underwent SVF therapy (**Table S6**), 2 patients exhibited complete recovery (disappearance of ulcer lesion, **Figure S7**) and 2 cases resulted in either major amputation or ulcer recurrence at the 1-year follow-up post-SVF therapy (**Figure 7B** and **7D**). All 4 patients had HOMA-IR within normal range (min-max 0.3-2.0, mean 1.4, **Table S7**). Based on the above results, we hypothesized that successful recovery at the 1-year follow up is correlated to the number of CD271^+^ progenitors initially injected. Thus, we retrospectively evaluated the potential impact of injected CD271^+^ progenitor number on efficacy of SVF therapy (**Figure 7C**). Then, we stratified patients based on recurrence or amputation outcome and revisited the composition of cell initially injected. Although pa 355 tient number in this analysis is very small, we observed that our primary outcome does not correlate with total SVF nor with the Lin^-^CD34^+^ cell number injected (**Figure 7E**). In particular, patient 2 who required amputation, had received the highest dose of total SVF cells and CD34^+^ cells compared to those of the patients who recovered, suggesting that high doses of CD34^+^ cells was insufficient for preventing amputation. However, both patients that completely recovered from ulcers without recurrence (patient 1 and 4) were injected with higher number of CD271^+^ cells (**Figure 7E**). As an example, Patient 1, who recovered from CLI, had approximately 200 times higher number of CD271^+^ progenitors injected than patient 3 who presented with recurrent ulceration (**Figure 7C** and **F**). The positive correlation between full recovery and CD271^+^ progenitors injected was striking especially when we compared the number of injected CD271^+^ progenitors per ulcer area (***n*** = 6 ulcer lesions, **Figure 7F**). Collectively, our findings from this sub-analysis of SVF therapy for CLI, strongly support the significance and positive therapeutic effects of CD271^+^ progenitors.

**Figure 7:**
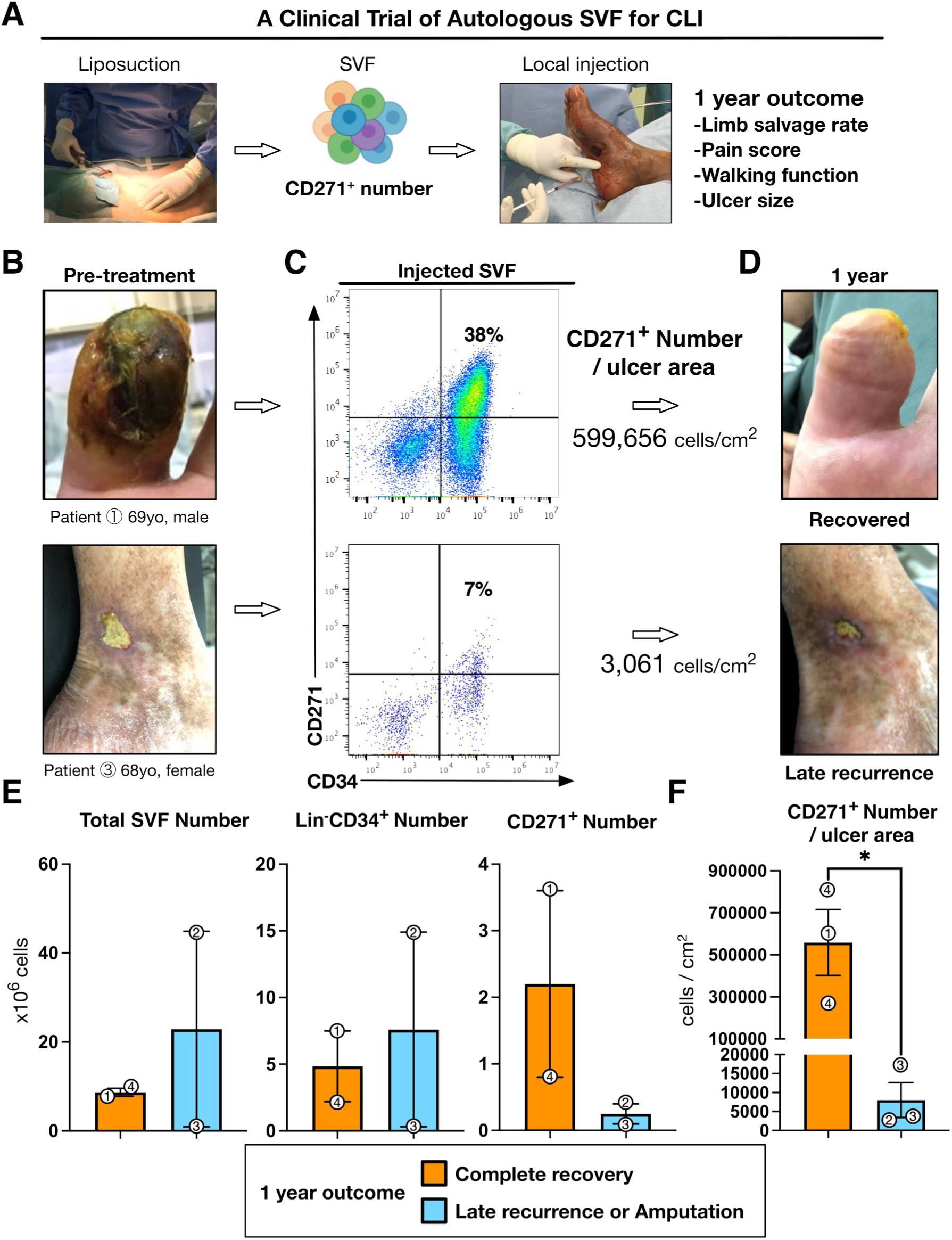
The number of AT-CD271^+^ progenitors injected was significantly higher in patients who completely recovered from CLI in SVF therapy. (A) Diagram of the study design of a clinical trial of autologous SVF transplantation for CLI. (B) Representative images of CLI lesions prior to autologous SVF transplantation. (B) Representative flow cytometric analysis of autologous SVF transplantation. (C) Representative images of CLI lesions acquired 12 months after autologous SVF transplantation. (E) Quantification of total SVF, Lin^-^CD34^+^, and CD271^+^ cell numbers injected to CLI patients (***n*** = 4) (F) Quantification of the number of CD271^+^ cells per ulcer area between patients (***n*** = 4 patients, ***n*** = 6 ulcer lesions). Numbers within data point indicates patient identification number. Data are represented as the mean ± SEM. \****P*** < 0.05. See also **Figure S7** and **Table S7**.

## DISCUSSION

Despite decades of research in therapeutic angiogenesis, the ideal cell population for grafting remains to be fully determined. This is evidenced by controversial effects on amputation prevention in 19 RCTs (837 patients) using 5 cell types^12^. In this study, we used unbiased single-cell transcriptomics to identify adipose tissue CD271^+^ progenitors as a human pro-angiogenic cell population with the strongest potential for efficacious therapeutics. Traditionally, the identification of cell populations suitable for cell therapy had depended on biased selection of markers, from which appropriate marker-positive cell populations were isolated for in-depth analysis *in vitro* and *in vivo*. This potential hit-and-miss strategy is time consuming and costly, and allow the analysis of only a limited number of markers.^49^ In this study, we took advantage of current single cell genomic approaches to simultaneously explore all known/detected surface markers. Based to these unbiased analyses of available scRNA-seq datasets, we found that AT-CD271^+^ progenitors were predominantly pro-angiogenic with less than 3% of antiangiogenic cells. We suggest that application of single-cell transcriptomics for cell therapy can greatly improve discovery of better cell markers for targeted therapies of interest. Furthermore, we took advantage of evaluating cell populations with opposing functions through scRNAseq analysis, i.e. therapeutic (pro-angiogenic) and harmful (anti-angiogenic) populations. Although an extra purification step to exclude anti-angiogenic cells was not necessary due to the extremely low rate of anti-angiogenic cells in CD271+ progenitors, sorting methodologies 386 to isolate therapeutic marker-positive away from harmful marker-negative cells could be strategic. For example, immune-modulatory cell therapy for graft-versus-host disease,^50, 51^ which is increasingly evaluated in clinical trials but with inconsistent outcomes, might benefit from such a strategy to avoid pro-inflammatory cell engraftment.

The relationship between CD271 expression and cell angiogenic capacity has been poorly established. There is a report demonstrating that CD271^+^ progenitors secreted angiogenic factors *in vitro*.^52^ However, limited angiogenesis was concomitantly observed in a mouse model of CD271^+^ cell injection for cartilage repair.^53^ Based on our present results, CD271^+^ progenitors could exhibit angiogenic capacity specifically in ischemic-type diseases. Interestingly, previous demonstrations that CD271^+^ cell/HUVEC coculture, but not conditioned medium, could enhance the tube formation of HUVECs, suggest the importance of cell-to-cell interactions between CD271^+^ progenitors and target cells. ^52, 53^ In our xenograft model, we observed CD271^+^ progenitors were surrounding capillaries and labeled by perfused lectin as with capillaries (**Figure 4B-C** and **Figure S3A**). This indicates the close proximities between “leaky” capillaries and engrafted cell grafts. Taken together, we conclude engraftment capability and cell-cell interactions between cell grafts and the regenerating (immature) capillaries would be important for CD271^+^ progenitors to exert angiogenic functions.

CD271, also known as LNGFR, is a transmembrane 404 receptor originally found to be a marker for neural lineage cells. CD271 is known to activate mTOR through cooperation with coreceptors including TRKB, also known as NTRK2 (neurotrophic receptor tyrosine kinase 2)^43, 54^ a coreceptor which we found to be abundantly expressed in the majority of CD271^+^ progenitors from human SVF cells. Data from single-cell transcriptomics, mutant CD271 mouse model, and mTOR silencing studies, all supported the participation of CD271/mTOR axis in the angiogenic capacity of CD271^+^ progenitors. However, our si-mTOR studies may reflect general blunted impacts since the mTOR pathway is a well-established major signaling hub for multiple factors ^55, 56^. Our results, nevertheless, highlight the importance of mTOR activity for evaluating ADSC quality. Future studies should assess whether neurotrophic factors may enhance CD271/TRKB/mTOR signaling in CD271^+^ progenitors and if neurotrophin pretreatment or controlled CD271 overexpression might be preconditioning strategies for enhancing angiogenic effects of cell grafts.

In this study, we observed proliferative and angiogenic capacity of AT-CD271^+^ progenitors. We found that angiogenic AT-CD271^+^ progenitor populations were reduced in patients with insulin resistance, although the causal relationship between AT-CD271^+^ progenitor decrease and insulin resistant is unclear from our study. Since blunted mTOR signaling in AT can lead to insulin resistance,^44, 57^ the CD271/mTOR axis in AT may have protective roles against insulin resistance. This may be important under obesogenic 422 stimuli (e.g., overfeeding), when AT dramatically increases its size and appropriate angiogenesis is essential in obesity to prevent hypoxia which triggers AT inflammation and subsequently insulin resistance.^58–61^ Future detailed investigations of the CD271/mTOR axis in AT will provide better understanding of insulin resistance pathophysiology and may reveal novel strategies to prevent metabolic syndrome.

Although preclinical studies suggested promising efficacy of cell therapy, clinical trials have shown inconsistent results. This might be explained by numerical or functional differences of cell grafts among CVD patients ^62, 63^. In this study, we demonstrated decreased cell number and functional impairment of CD271^+^ progenitors in insulin resistant donors. Given the higher incidence of metabolic disorders in CVD patients, our findings have great implications for effective cell therapy in CVD. It is critical to overcome the reduction in cell number and the functional impairment of CD271^+^ progenitors in insulin resistant donors if efficient CD271^+^ cell therapy is to be developed. We and others have previously reported altered gene expression and impaired angiogenic effects of ADSCs derived from type 2 diabetic patients compared to ADSCs from nondiabetics.^64, 65^ In this study, we found that HOMA-IR can be used as a surrogate marker to predict quantity and quality of CD271^+^ progenitors in AT. Therefore, HOMA-IR evaluations may mitigate ineffective therapeutic interventions by permitting screening of appropriate donors for therapeutic angiogenesis, not only for autologous therapy, but also for allograft (cell bank) development.

Our single cell transcriptomics from multiple stem cell sources captured AT as a better cell source for angiogenic cell populations than current BM mononuclear cells. Furthermore, we showed CD34^+^ selection did not increase the quantity of appropriate angiogenic cell population when compared to unsorted SVF (**Figure 2H**). Here, we propose that targeted CD271 selection is a novel strategy for preparing cell graft with CD271^+^ progenitor therapy expected to improve cell therapeutic outcomes. Although CD271^+^ progenitors showed more angiogenic gene profiles over the other candidate cell populations we screened (ABCB11 and VCAM1) or previously reported (BM-CD34^+^, CD133^+^, and CD31^+^), we did not compare angiogenic capacity directly in this study. Further studies to compare CD271^+^ progenitors and other potential subpopulations *in vitro* and *in vivo* would be important to develop the most suitable strategy for therapeutic angiogenesis. More importantly, as demonstrated in this study (**Figure 6** and **7**), the feasibility/accessibility of these cell populations in human donors should be tested to successfully translate our findings into clinical trials.

Our retrospective analysis of the SVF therapy trial for terminal CLI ^47^ supports a positive correlation between complete recovery and high numbers of injected CD271^+^ progenitors, while amputated/recurrence of CLI was associated with low numbers. Importantly, we determined that approximately 300,000 to 500,000 AT-CD271^+^ progenitors per lesion area (CD271+ cells / ulcer cm2) as the dose that seems 458 to be necessary for favorable outcome.

Fortunately, this is a dose which, in our experience, should be readily feasible to prepare from CVD patients. Future studies and clinical trials focusing on CD271^+^ progenitor therapy in larger cohorts will provide the necessary clinical evidence for efficacy and dosing requirements.

In summary, we applied single-cell transcriptomics for cell therapy purposes against CLI and identified a human stem cell population which is readily available and possesses the most angiogenic gene profile. CD271^+^ progenitors can be readily harvested from human subcutaneous AT and exhibit promising angiogenic capacities, including long-term engraftment and augmentation of host tissue cell proliferation in cell therapy. In CVD patients with systemic insulin resistance, the number and angiogenic capacity of CD271^+^ progenitors were significantly decreased. CD271^+^ progenitors obtained from insulin sensitive donors, in contrast, demonstrated the highest *in vivo* angiogenic capacity and consistently rescued ischemic limb. Our data highlight the potential screening benefits using single-cell technologies and evident benefits of CD271^+^ progenitor transplantation for improving current therapeutic angiogenesis.

### Study Limitations

In this study, the correlation between the expression levels of CD271 mRNA and protein was not directly analyzed. Although we confirmed higher mRNA expression of CD271 in sorted CD271^+^ progenitors (data not shown), further studies using single-cell multiomics technologies, including CITE-seq,^66^ would be helpful to validate our results.

In mouse ADSC experiments, we observed that a mutation of the CD271-encoding gene decreased angiogenic factor expression and impaired *in vivo* angiogenic capacity in unsorted mouse ADSCs. We did not sort CD271^+^ cells from CD271^Mut^ mice because of difficulties in detecting mutated CD271 with currently available monoclonal antibodies.

Finally, we could not definitely determine whether injected CD271^+^ progenitors transdifferentiate into endothelial cells or not. Although expression of human CD31 was observed in only a small proportion of engrafted CD271^+^ progenitors, the histological assessment demonstrated that they were morphologically similar to stromal or perivascular cells than to endothelial cells. Single-cell analysis of regenerating tissues after cell therapy might provide more detailed information about the transcriptomic dynamics of injected CD271^+^ progenitors.

## ACKNOWLEDGEMENTS

We would like to acknowledge the support and collaborations of Professor Toyoaki Murohara, Dr. Rei Shibata, Dr. Yuuki Shimizu, and TACT-ADRC multicenter trial Group regarding SVF therapy trial. Additionally, we appreciate Professor Yoh Takuwa and Dr. Kazuaki Yoshioka for the assistance in Laser Doppler analysis for blood flow quantification. Furthermore, we thank Dr. Jun Kinoshita, Dr. Katsunobu Oyama, Dr. Yukio Fushida, and the other operating staff of Kanazawa University Hospital for providing adipose tissue samples to this study. Finally, we appreciate the study participants and their families. Diagrams in Figures were created with BioRender.com.

## FUNDING

This study has been supported by Japan Society for the Promotion of Science (JSPS) KAKENHI (Tokyo, Japan) Grant Number JP16H06828. OI is supported by Japanese Heart Foundation Research Abroad Award 2021. JSG is supported by grants from the American Heart Association (18CDA34080527), the NIH (R21OD031907), a CCHMC Trustee Award, a CCHMC Center for Pediatric Genomics Award and a CCHMC Center for Mendelian Genomics & Therapeutics Award. The content is solely the responsibility of the authors and does not necessarily represent the official views of the National Institutes of Health.

## AUTHOR CONTRIBUTIONS

Conceptualization: O.I., V.H., J.S.G., S.U., M.T.

Methodology: O.I., C.G., D.H., K.Y., Y.T., A.N., H.M., J.S.G., S.U. Validation: O.I., C.G., D.H., K.Y., Y.T., A.N., H.M., J.S.G., S.U. Formal Analysis: O.I., A.N., D.H., K.Y., H.W.L., J.S.G., S.U.

Investigation: O.I., C.G., D.H., K.Y., Y.T., A.N., H.O., S.T., H.M., H.W.L., J.S.G., S.U. Resources: K.I., H.T.

Data Curation: O.I., H.W.L., V.H., J.S.G., S.U.

Writing – Original Draft: O.I., C.G., V.H., J.S.G., S.U., M.T.

Writing – Review & Editing: O.I., C.G., D.H., K.Y., Y.T., A.N., H.O., S.T., K.I., H.T., H.M., H.W.L., V.H., J.S.G., S.U., M.T.

Visualization: O.I., H.W.L., V.H., J.S.G., S.U. Supervision: J.S.G., S.U., M.T.

Project Administration: J.S.G., S.U., M.T. Funding Acquisition: J.S.G., S.U., M.T.

## COMPETING INTERESTS

The authors declare no competing interests.

## METHODS

### RESOURCE AVAILABILITY

#### Lead contact

Further information and requests for resources and reagents will be fulfilled by the lead contact, Masayuki Takamura

#### Materials availability

None of unique reagents were generated in this study.

#### Data and code availability

Single-cell RNA-seq datasets analyzed in this study are available at NCBI Gene Expression Omnibus (GEO). GEO accession numbers are listed in **Table S1**. Any information and Seurat analysis files required to reanalyze the data reported in this study is available from the Lead Contact upon request.

## EXPERIMENTAL MODEL AND SUBJECT DETAILS

### Study Design and Patient Population

In the study of patient-derived AT and SVF, eligible patients were adult males or females with CVD who underwent elective cardiac surgery, including coronary artery bypass grafting (CABG) and cardiac valve replacement/plastic surgeries. Major exclusion criteria were acute emergency diseases, including acute myocardial infarction, active and acute infectious disease, uncontrolled diabetes mellitus (HbA1c > 8.0%), hypo/hyperinsulinemia (< 2.2 or > 13.0 μU/mL), systemic inflammatory disease (autoimmune disease), and history of irradiation therapy at tissue harvest sites. Thirty-eight patients were enrolled between April 2017 and February 2022. This study was registered in the University Hospital Medical Information Network Center (UMIN, Tokyo, Japan) Clinical Trials Registration System as UMIN000020764. After enrollment with written informed consent, clinical and laboratory variables were collected preoperatively, and all data were blinded for the researchers who performed further experiments.

The detailed protocol and eligibility for the clinical trial of SVF therapy for CLI are published,^47, 67^ Briefly, this was a multicenter clinical trial (UMIN000010143) to investigate the feasibility and safety of SVF injection for CLI patients with no option for treatment^47^.

Autologous SVF from liposuction was transplanted to the ulcer lesions via intra-muscular injections. The safety, feasibility, and the efficacy including hard outcomes, including survival rate and amputation free rate in 6 months, of SVF therapy were investigated. In this study, we sub-analyzed 4 patients (**Table S6**) on whom we performed cell therapy in our institution (Kanazawa University) with 1-year follow-up data. On the remaining samples after SVF injection, we analyzed their viability and surface antigen expressions.

The studies using human samples subjects was performed according to Declaration of Helsinki principles and approved by the Research Ethics Board of the University of Kanazawa. Written informed consent was obtained from all patients prior to inclusion in the study. All animal experiments were approved by the Institution for Experimental Animals of the Kanazawa University Advanced Science Research Center.

### Mice

All animal protocols were approved and performed according to the Guide for the Care and Use of Laboratory Animals of Kanazawa University, which strictly conforms to the Guide for the Care and Use of Laboratory Animals published by the US National Institutes of Health (NIH, Bethesda, MD, USA). Athymic nude mice for xenograft experiments were purchased from the Charles River Laboratories. Heterozygous Ngfr^tm1j^^ae^^42^ were purchased from the Jackson Laboratory (Strain #002213) and crossed two-or three-times with C57BL/6J females before generating homozygous Ngfr^tm1Jae^ (CD271^Mut^) mice. Homozygous Ngfr^tm1Jae^ and controls mice were used at 6–8-week-old) before they develop peripheral nervous denervation and cutaneous ulcers (16 weeks old).

### Cell Therapy Model of Limb Ischemia

6-to 8-week-old athymic nude mice (Jackson Laboratory) were subjected to hindlimb ischemia. Mice were anesthetized by intraperitoneal injection of sodium pentobarbital (40 mg/kg). The proximal portion of the left femoral artery was resected. After 24 h, 0.5 million human ADSCs at passage 4-6 were prepared for injection. In brief, ADSCs were harvested by trypsinization, labeled with PKH26 cell linker dye (2μL/mL, 3 minutes in room temperature, Sigma), and resuspended in PBS (5.0 × 10^5^ cells in 60 μl with 27 gauge syringes). Labeled ADSCs were immediately injected intra-muscularly at three different sites (gastrocnemius, gracilis, and quadriceps muscles, 20 μl per injection). Sham-operated mice were injected with PBS and used as a control group. At day 7, 14, and 35 after cell therapy, mice were anesthetized and subjected to laser Doppler analysis for blood flow, followed by harvest of gastrocnemius muscles.

## METHOD DETAILS

### Single-Cell Transcriptome Datasets and Analysis

We used scRNA-seq datasets of AT,^26, 68, 69^ BM,^24, 70^ and UCB.^25, 71^ Gene Expression Omnibus (GEO) accession numbers of revisited datasets are listed in **Table S1**. For quality control, cells expressing < 200 genes and with a mitochondrial content > 5% were removed. For batch integration of multiple scRNAseq datasets, Harmony was used to remove the batch effect.^72^ After data normalization, dimension reduction (tSNE) and clustering with using a Louvain algorism were performed. Marker gene analysis and differential expression (DE) analysis were performed using non-parametric approach, Venice^73^ or DESeq2. The Gene Set Enrichment Analysis (GSEA) was performed to list GO Biological Process terms.^74^ Cluster annotation was determined according to marker genes for clusters identified by non-parametric approach, Venice and collation with overlapping genes from the Human Cell Atlas and previously published scRNA-seq reports.^19, 21^ These analyses were performed in BioTuring Browser 3.^75^

### Adipose Tissue and SVF Processing

Unnecessary subcutaneous adipose tissues (1-2 gram) discarded during making surgical incision were obtained. SVF cells were isolated from adipose tissue using a procedure modified from Zuk et al.^39^ Briefly, adipose tissue was digested by Liberase TM Research Grade collagenase solution (0.4 units/mL; Sigma-Aldrich) for 30 min using gentleMACS Dissociator (“ATDK” program, Miltenyi Biotec). Then, the digested solution was filtered through a 100-µm cell strainer (BD Biosciences) to remove undigested connective tissue debris. After centrifugation (400g, 10 minutes at room temperature), the supernatant containing mature adipocytes was discarded. The pellet was lysed to deplete erythrocytes with BD Pharm Lyse (2mL for 1 minute at room temperature, BD Biosciences) and washed twice with phosphate-buffered saline (PBS) with 2.0% fetal bovine serum (FBS). The final pellet was resuspended in 150 μL of autoMACS Running buffer (Miltenyi) and counted with a hemocytometer in triplicate.

### Flow Cytometry and Sorting

SVF cells were separated into CD45^+^ and CD45^-^ cells using microbeads and autoMACS pro separator (Miltenyi) according to the manufacturer’s instructions. CD45^-^ cells were incubated with Fc Block (BD Biosciences) prior to staining. Then, cells were labeled with following antibodies: human CD31-FITC (BD Biosciences, clone WM59, used in 1:20 dilution), CD271-APC (Miltenyi, cloneME20.4-1.H4, 1:20), and CD34-peridinin chlorophyll protein (PerCP) (BD Biosciences, clone 8G12, 1:20) for 15 min at 4°C. Then, cells were washed twice in PBS with 2.0% FBS. Flow cytometric data were acquired by using Accuri C6 (BD Biosciences). Before immunophenotyping, the instrument performance was validated by using BD Accuri Spherotech Validation Beads (BD Biosciences). A total of 30,000-100,000 events were acquired at slow flow rate. Acquired data were analyzed using FlowJo software X 10.0.7 (Tree Star Inc.). Isotype-matched negative controls or fluorescence-minus controls were used to define the threshold for each specific signal and establish the appropriate gate for positive cells.

### Cell Culture

The CD45^-^ stromal cells that remained from flow cytometric analysis were seeded at a density of 1.0 × 10^4^ cells/cm^2^. Cells were cultured in standard conditions (5% carbon dioxide, 37°C) in DMEM/F12 (Thermo Fisher Scientific) supplemented with 10% FBS and 1% antibiotic-antimycotic (Thermo Fisher Scientific) solution. At 24 h after seeding, non-adherent cells were removed by washing the plates, and then medium was changed every three days. ADSCs were harvested at sub-confluency with trypsin (0.05% trypsin-EDTA; Thermo Fisher Scientific). Cells at the second passage and thereafter were replated at a density of 3,000 cells/cm2.

### RT-qPCR

Total RNA was isolated from ADSCs using RNeasy Mini Kit (Qiagen). Total RNA (100 ng) was used to generate cDNA using the TaqMan Universal Master Mix (Applied Biosystems). Quantitative real-time polymerase chain reaction (qRT-PCR) analysis was performed on an ABI Prism 7300 sequence detection system (Applied Biosystems). The TaqMan probes (Applied Biosystems) used in this study were listed in **Table S7**. Relative expression was assessed by comparative CT method, correcting for amplification efficiency of the primers, and performed in duplicate.

### Colony-Forming Unit Assay

Cultured AT-CD271^+^ or CD271^-^ progenitors at passage 4 were used for colony-forming unit (CFU) assays. Cells were detached and plated at a density of 20 cells/cm^2^ in 10 cm dishes. Half the medium was changed at day 7 after plating. Adherent cell clusters of greater than 50 cells were counted at day 14 with Giemsa staining. The experiments were performed in triplicate for each sample.

### Flow Cytometry of ADSC treated limb

At 14 days after cell therapy in xenograft model, gastrocnemius muscle was harvested and digested using gentleMACS (Miltenyi) with “m_muscle_01” program and Liberase TM solution. Single cells from digested limb were filtered through 40μm cell strainer and labeled with immunofluorescence-conjugated antibodies; mice CD31-PerCP-Cy^5^.^5^ (BD Bioscience, clone MEC 13.3, 1:20), human CD31-FITC (BD Biosciences, clone WM59), human CD271-APC (Miltenyi, cloneME20.4-1.H4, 1:20), human CD90-APC (BD Biosciences, clone 5E10), and human CD105-PerCP-Cy^5.5^ (BD Biosciences, clone 266, 1:20) for 15 min at 4°C. Flow cytometric data was acquired by similar methods described above in *Flow Cytometry and Sorting* section.

### Blood Flow Measurement

Blood flow was assessed using Laser Doppler blood flow imager (OZ-2 LCB, OMEGAWAVE, Japan) at day 1, 3, and 14 after induction of limb ischemia. Before scanning, mice were placed on a heating plate at 37°C for 15 min, followed by anesthetization using intraperitoneal injection of sodium pentobarbital (20 mg/kg). The signal intensities basically reflecting laser scattering by red blood cells within circulation were obtained as blood flow value. The mean blood flow value was acquired in region of interest (ROI) capturing the whole mouse lower limb. The ratio of blood flow values of the ischemic (left) limb to the healthy (right) limb were calculated as blood flow recovery to avoid data variations caused by ambient light and temperature.^8, 9^

### In Vivo Proliferation Assay

EdU solution (5mg/2mL) was administered via intra-peritoneal injection at day 6 after ADSC therapy. At day 7, the gastrocnemius muscles were harvested and processed to frozen sections. EdU positive nucleus were detected using EdU *in vivo* imaging kits (BCK488-IV-IM, baseclick) according to the manufacturer’s instruction. The number of EdU^+^ and DAPI^+^ nucleus was counted in frozen sections in duplicate.

### Immunohistochemistry

Tissue vascularization and ADSCs engraftment were evaluated at day 14 and 35. Mice were anesthetized and injected with 50 μg of FITC-conjugated isolectin B4 (FL-1201, Vector Laboratories) via tail vein. After 10 min, mice were perfused with PBS for 3 min to remove unbound isolectin B4 in the blood vessels, and perfusion-fixed with 4% paraformaldehyde. The gastrocnemius muscles were removed, fixed with 4% paraformaldehyde at 4°C overnight, and incubated in 30% sucrose solution for 24 hours. Frozen sections of the tissue were obtained at 10 μm thickness and counterstained with DAPI. Capillary density and engraftment of injected ADSCs were analyzed by counting FITC and PKH positive cells, respectively, in five randomly selected fields per section. The experiments were performed in duplicate or triplicate for each human ADSCs sample.

### Statistics

Continuous data are expressed as the mean ± SEM, and statistical analysis was performed with Prism 9 software (GraphPad, USA). Group comparisons were performed using paired or unpaired *t* tests or one-way analysis of variance (ANOVA). Correlation analysis was performed using Pearson correlation for normally distributed data (Shapiro–Wilk normality test) or Spearman correlation for nonnormally distributed data. Statistical significance was defined as \****P*** < 0.05, \*\****P*** < 0.01, \*\*\****P*** < 0.005 and \*\*\*\****P*** < 0.001. All reported statistical tests were two-tailed.

## Notes

### Competing Interest Statement

The authors have declared no competing interest.

### Summary of Updates

This revision includes modifications in Graphical Abstract and text formatting.

