## Supplemental Files for "Single cell transcriptomics identifies adipose tissue CD271+ progenitors for enhanced angiogenesis in limb ischemia"

### Figure S1

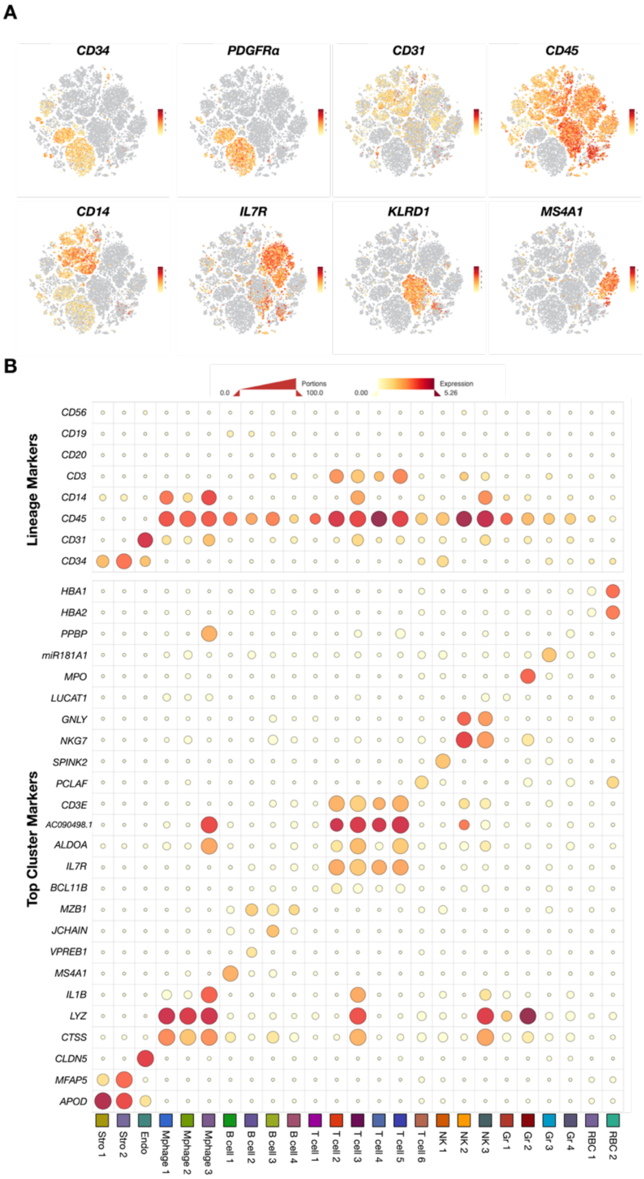


**Figure S1: Cluster annotations of lineage-negative cells from multiple stem cell sources**
(A) tSNE map and scatter plots of marker genes in lineage-negative cells from BM, AT, and UCB (***n*** = 2/tissue).
(B) Heatmap presentation of marker gene expression for clusters. The size of the dots reflects the proportion of cells that express the gene. The color of the dots shows the gene expression levels.

**Figure S2**

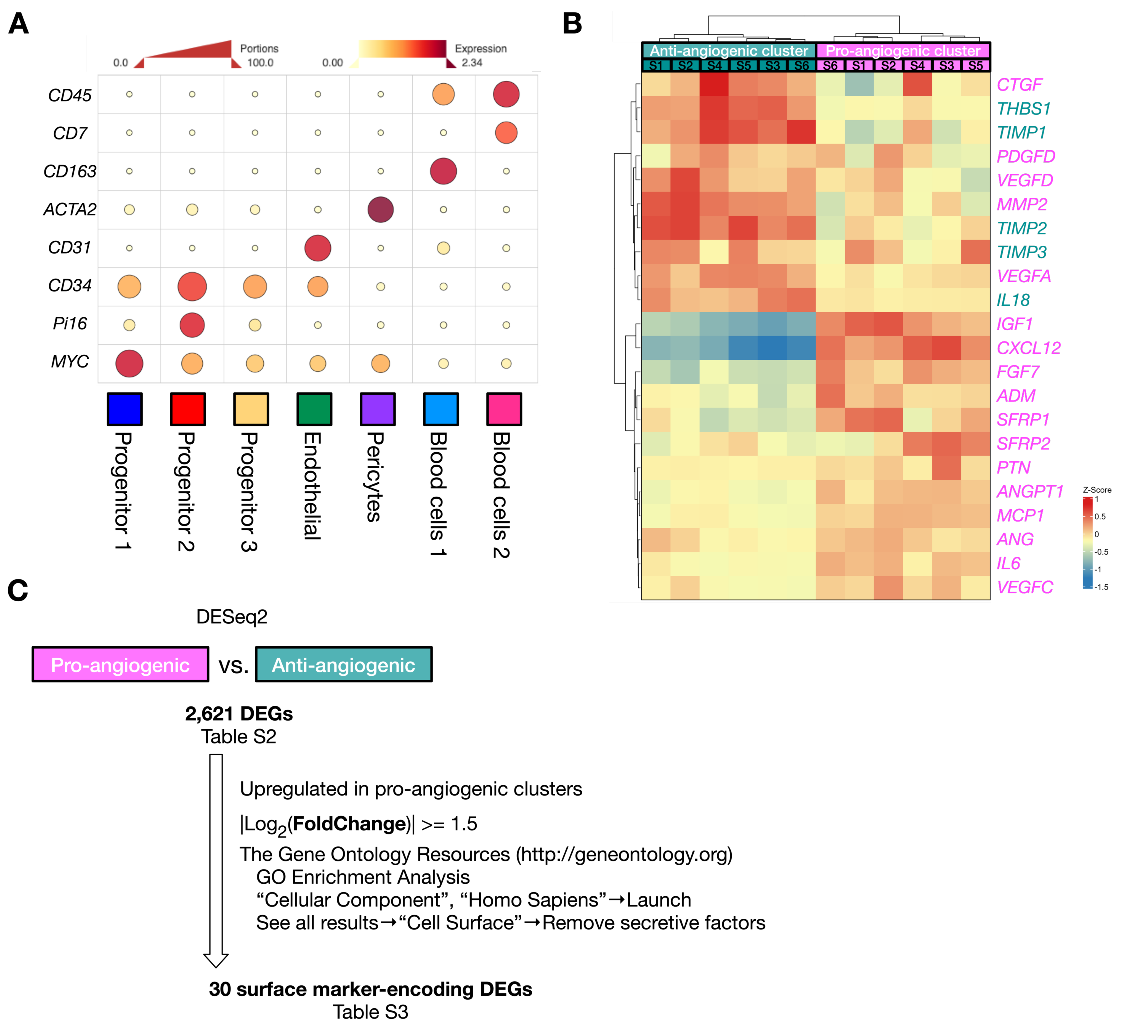


**Figure S2: Filtering strategy of angiogenic cell population and comparison with previously reported angiogenic cell populations**

(A) Heatmap presentation of z-score expression levels of marker genes defined by Marker Analysis across the clusters and representative known genes in human SVF scRNA-seq analysis (***n*** = 6).

(B) Heatmap of 22 pro/anti-angiogenic factor genes between pro/anti-angiogenic clusters. The color of the bar shows the gene expression levels quantified by the Z-score (***n*** = 6).

(C) Diagram to summarize the filtering analysis to screen 30 surface marker genes specifically expressed in the pro-angiogenic cluster.

**Figure S3**

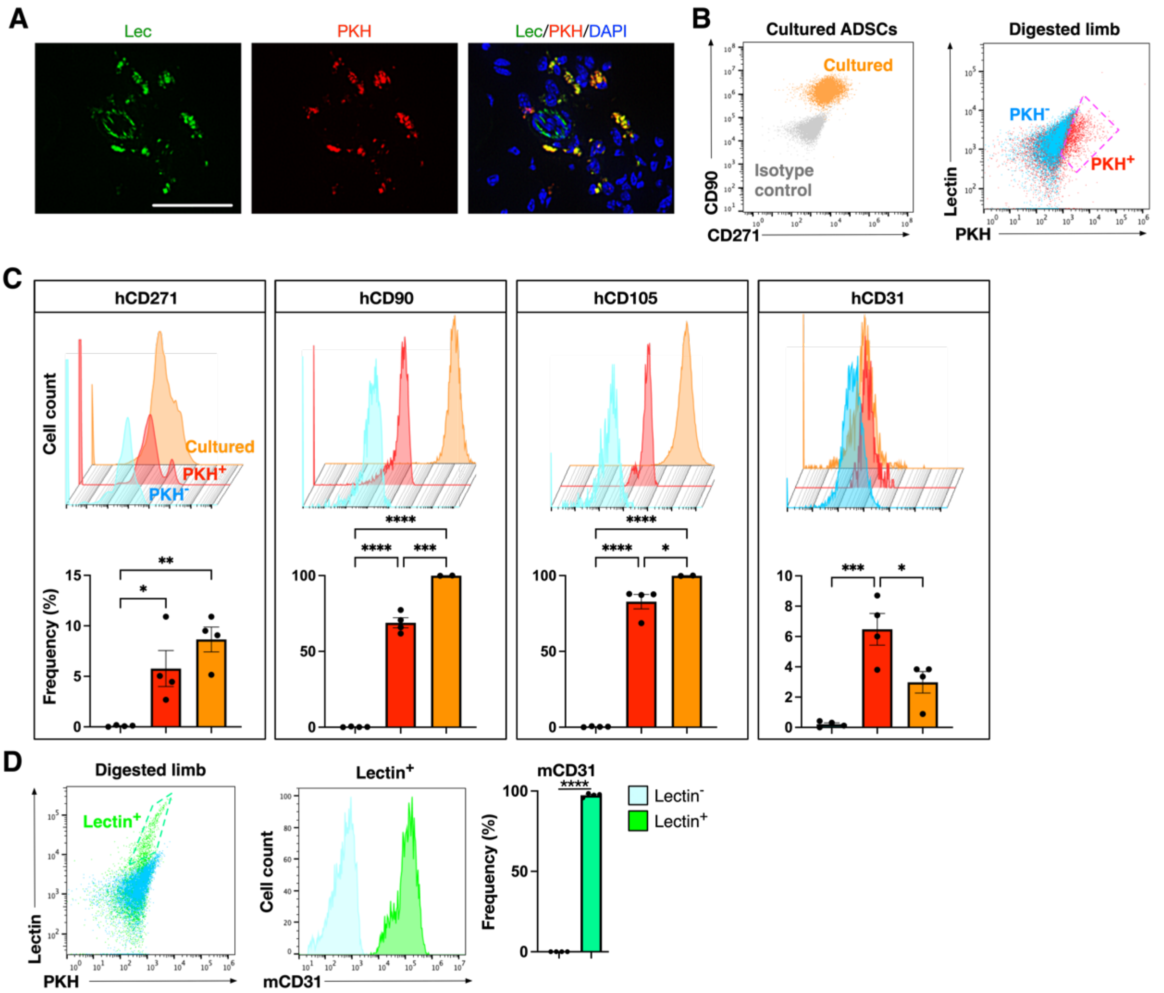


**Figure S3: Localization and immunophenotypic characterization of Lectin$^+$ capillaries and PKH$^+$ ADSCs in cell therapy experiments**

(A) Representative IHC images of Lectin$^+$ and PKH$^+$ cells in mice limb on day 14 after cell transplantation.

(B) Representative flowcytometric plots of cultured ADSCs and PKH$^+$ ADSCs after cell therapy.

(C) Comparative analysis of human ADSC/endothelial marker (hCD31) expression between cultured ADSCs, PKH$^+$ ADSCs, and PKH$^-$ cells after cell therapy (***n*** = 2-4).

(D) Representative flowcytometric plots and comparative analysis of mice endothelial marker (mCD31) expression between Lec$^+$ and Lec$^-$ cells after cell therapy (***n*** = 4). Data are represented as the mean ± SEM. *$P < 0.05$, **$P < 0.01$, ***$P < 0.005$ and ****$P < 0.001$.

**Figure S4**

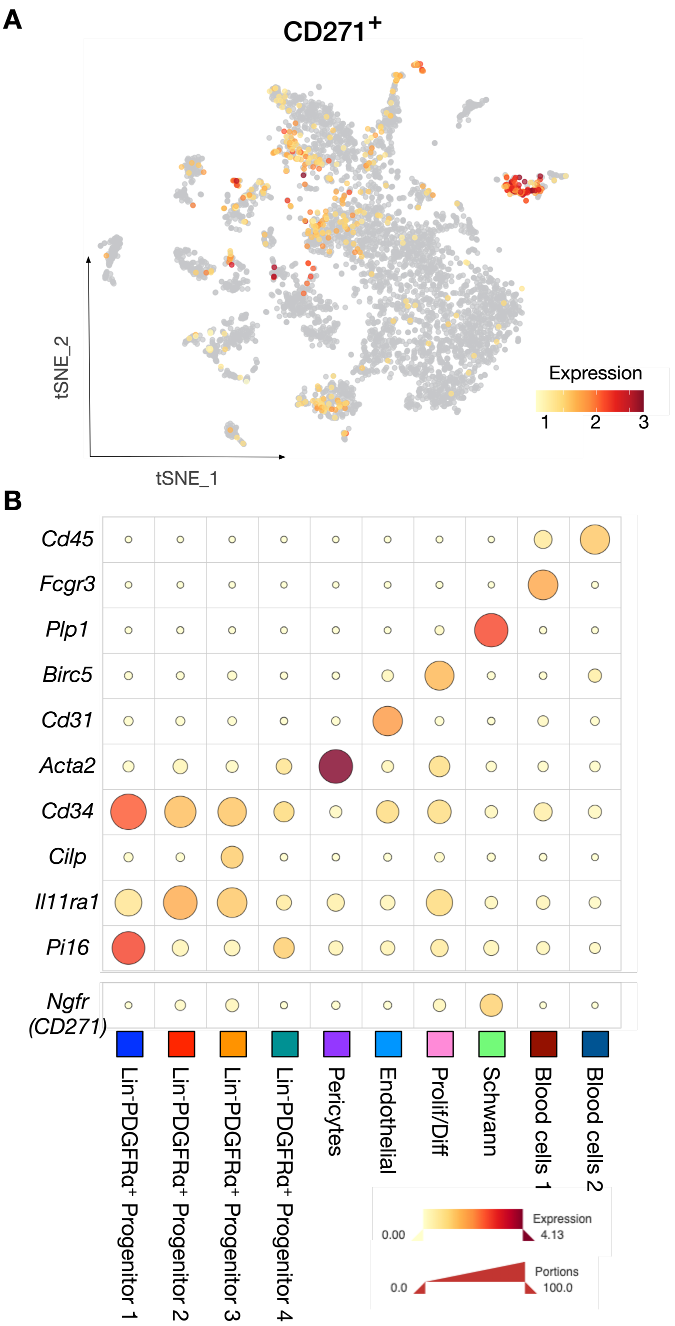


**Figure S4: Characterization of CD271$^+$ cells in mice SVF.**
(A) The distribution of CD271$^+$ cells in tSNE plot of mice SVF.
(B) Heatmap of *Ngfr* (CD271) and marker gene expression for clusters in mouse SVF. The size of the dots reflects the proportion of cells that express the gene. The color of the dots shows the normalized gene expression levels.

**Figure S5**

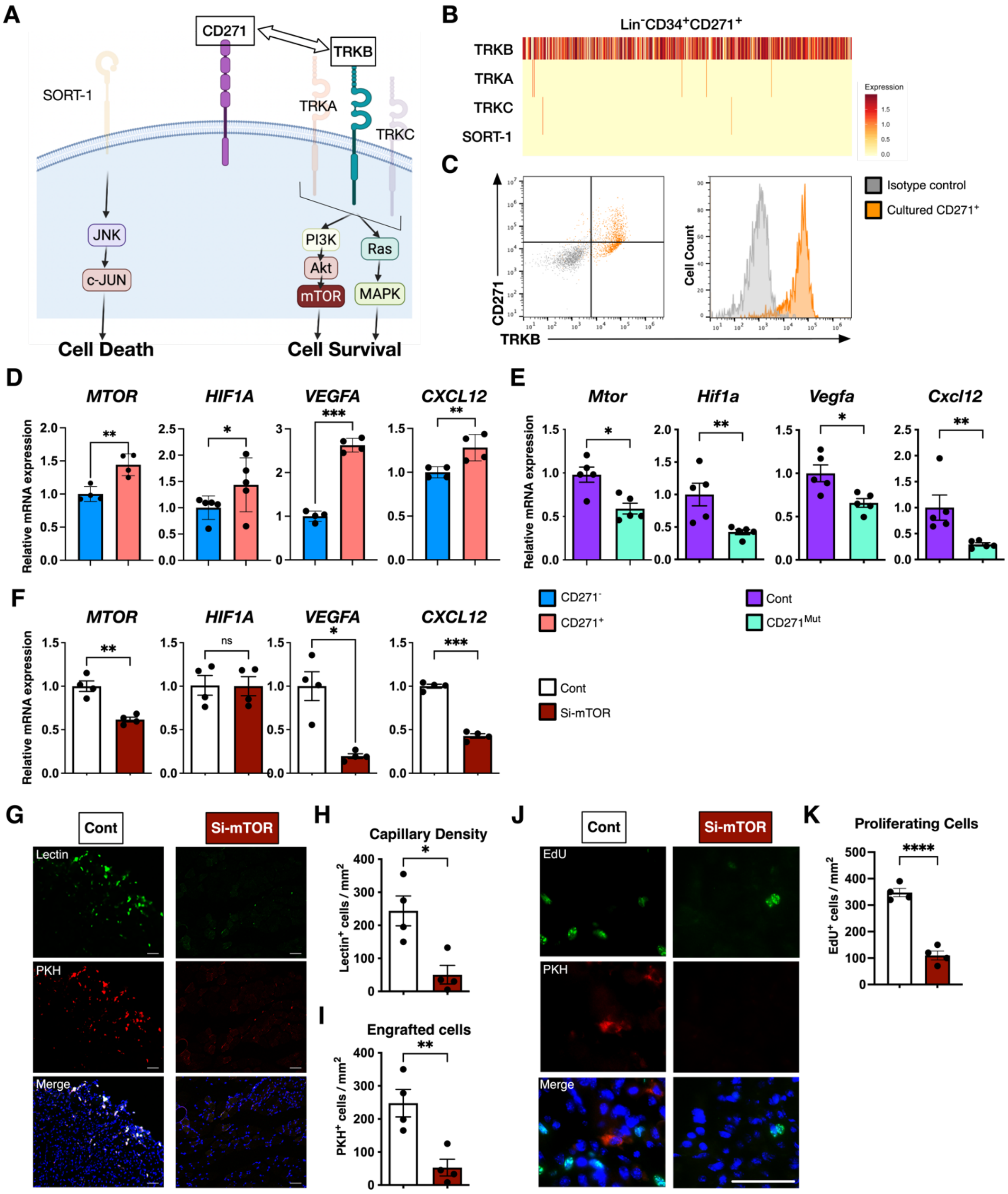
A
CD271
TRKB
SORT-1
TRKA
TRKC
JNK
c-JUN
PI3K
Akt
mTOR
Ras
MAPK
Cell Death
Cell Survival
B
Lin⁻CD34⁺CD271⁺
TRKB
TRKA
TRKC
SORT-1
Expression
C
CD271
TRKB
Cell Count
Isotype control
Cultured CD271⁺
D
MTOR
HIF1A
VEGFA
CXCL12
Relative mRNA expression
E
Mtor
Hif1a
Vegfa
Cxcl12
Relative mRNA expression
CD271⁻
CD271⁺
Cont
CD271Mut
F
MTOR
HIF1A
VEGFA
CXCL12
Relative mRNA expression
ns
Cont
Si-mTOR
G
Cont
Si-mTOR
Lectin
PKH
Merge
H
Capillary Density
Lectin⁺ cells / mm²
I
Engrafted cells
PKH⁺ cells / mm²
J
Cont
Si-mTOR
EdU
PKH
Merge
K
Proliferating Cells
EdU⁺ cells / mm²

**Figure S5: CD271/TrkB-mTOR axis has a role in angiogenic potential of human ADSCs**

(A) Diagram illustrating coreceptors of CD271 and corresponding cell signaling pathways.

(B) Single-cell heatmap presentation of the gene expression levels of coreceptors in $Lin^-$ $CD34^+CD271^+$ cells. The color of the heatmap represents gene expression levels. Each bar shows gene expression of a single cell of 741 $Lin^-CD34^+CD271^+$ cells (***n*** = 6).

(C) Representative flowcytometric plots of CD271 and TrkB expression in $CD271^+$ ADSCs.

(D) RT-qPCR of *MTOR, HIF1A, VEGFA, and CXCL12* expression between $CD271^+$ and $CD271^-$ progenitors (***n*** = 4). The mean value of $CD271^-$ progenitors is set as 1.

(E) RT-qPCR of *Mtor, Hif1a, Vegfa, Cxcl12* expression between mice Control-ADSCs and $CD271^{mut}$ ADSCs (***n*** = 5). The mean value of Control-ADSCs is set as 1.

(F) RT-qPCR of *MTOR, HIF1A, VEGFA, and CXCL12* expression between human ADSCs treated with siRNA for mTOR (Si-mTOR)-and control (Control, ***n*** = 4).

(G) Representative IHC images of $lectin^+$ capillaries and $PKH^+$ engrafted human ADSCs at day 7 after cell transplantation of Control-ADSCs and Si-mTOR ADSCs. Bar = 100 μm.

(H and I) Quantitative analysis of (H) capillary density and (I) engrafted ADSCs in ischemic limbs on day 7 (***n*** = 4).

(J) Representative IHC images of $EdU^+$ proliferating cells at day 7 after cell transplantation of Control-ADSCs and Si-mTOR ADSCs. Bar = 100 μm.

(K) Quantitative analysis of proliferating cells in ischemic limbs on day 7 (***n*** = 4). The mean value of Si-Control ADSCs is set as 1. Data are represented as the mean ± SEM. *$P < 0.05$, **$P < 0.01$, ***$P < 0.005$ and ****$P < 0.001$.

**Figure S6**

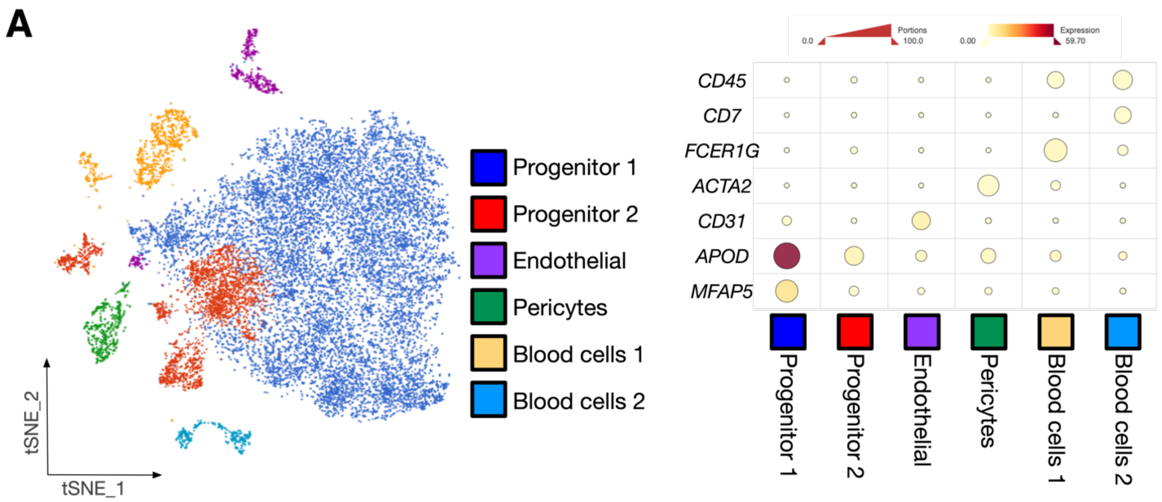


**Figure S6: Determination of a target value of capillary density to achieve complete recovery of blood flow**
(A) tSNE visualization and heatmap presentation of marker gene expression for 6 clusters in SVF from insulin sensitive and resistant donors. The size of the dots reflects the proportion of cells that express the gene. The color of the dots shows the normalized gene expression levels.

**Figure S7**

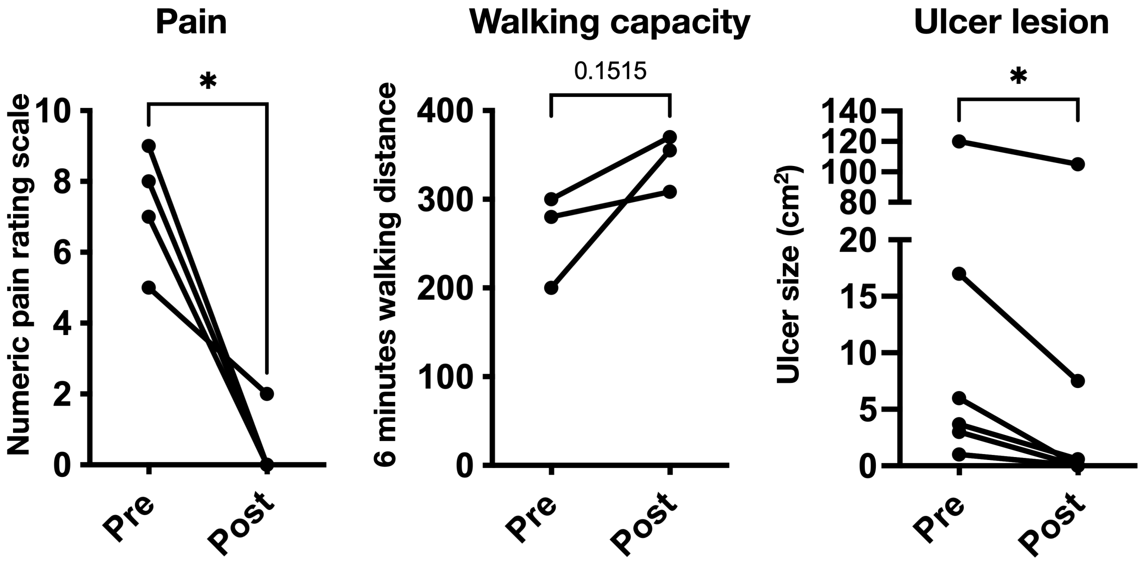


**Figure S7: Outcomes of SVF therapy at 1-year follow-up**

Comparative analysis of numerical pain scale score (***n*** = 4), 6 minutes walking distance (***n*** = 3, one patient failed the measurement because of limb amputation), and total ulcer lesion area (***n*** = 6) between pre- and post- SVF therapy were displayed. Data are represented as the mean ± SEM. ****P*** < 0.05.

**SUPPLEMENTAL TABLES**

**Table S1:** Datasets used in scRNA-seq analysis in **Figures 1, 2, 3, 5, and 6**.

**Table S2:** Gene set of secreted pro-/anti-angiogenic factors used in **Figures 1 and 2**.

**Table S3:** Differentially expressed genes between pro- and anti-angiogenic clusters in **Figure 2**.

**Table S4:** Differentially expressed genes encoding cell surface protein between pro- and anti-angiogenic clusters (Pro > Anti) in **Figure 2**.

**Table S5:** Patient characteristics in **Figure 6**.

**Table S6:** Pathway enrichment analysis between $CD271^+$ cells from insulin sensitive and resistant donors (Sensitive > Resistant) in **Figure 6**.

**Table S7:** Baseline characteristics of 4 CLI patients receiving SVF therapy in **Figure 7**.

**Table S8:** Taqman probes used in RT-qPCR assays.
